# Efficient coding, channel capacity and the emergence of retinal mosaics

**DOI:** 10.1101/2022.08.29.505726

**Authors:** Na Young Jun, Greg D. Field, John M. Pearson

## Abstract

Among the most striking features of retinal organization is the grouping of its output neurons, the retinal ganglion cells (RGCs), into a diversity of functional types. Each of these types exhibits a mosaic-like organization of receptive fields (RFs) that tiles the retina and visual space. Previous work has shown that many features of RGC organization, including the existence of ON and OFF cell types, the structure of spatial RFs, and their relative arrangement, can be predicted on the basis of efficient coding theory. This theory posits that the nervous system is organized to maximize information in its encoding of stimuli while minimizing metabolic costs. Here, we use efficient coding theory to present a comprehensive account of mosaic organization in the case of natural videos as the retinal channel capacity—the number of simulated RGCs available for encoding—is varied. We show that mosaic density increases with channel capacity up to a series of critical points at which, surprisingly, new cell types emerge. Each successive cell type focuses on increasingly high temporal frequencies and integrates signals over large spatial areas. In addition, we show theoretically and in simulation that a transition from mosaic alignment to anti-alignment across pairs of cell types is observed with increasing output noise and decreasing input noise. Together, these results offer a unified perspective on the relationship between retinal mosaics, efficient coding, and channel capacity that can help to explain the stunning functional diversity of retinal cell types.

## 1 Introduction

The retina is one of the most intensely studied neural circuits, yet we still lack a computational understanding of its organization in relation to its function. At a structural level, the retina forms a three-layer circuit, with its primary feedforward pathway consisting of photoreceptors to bipolar cells to retinal ganglion cells (RGCs), the axons of which form the optic nerve [1]. RGCs can be divided into 30-50 functionally distinct cell types (depending on species) with each cell responsive to a localized area of visual space (its receptive field (RF)), and the collection of RFs for each type tiling space to form a “mosaic” [2, 3, 4, 5]. Each mosaic represents the extraction of a specific type of information across the visual scene by a particular cell type, with different mosaics responding to light increments or decrements (ON and OFF cells), high or low spatial and temporal frequencies, color, motion, and a host of other features. While much is known about the response properties of each RGC type, the computational principles that drive RGC diversity remain unclear.

Efficient coding theory has proven one of the most powerful ideas for understanding retinal organization and sensory processing. Efficient coding posits that the nervous system attempts to encode sensory input by minimizing redundancy subject to biological costs and constraints [6, 7]. As more commonly formulated, it seeks to maximize the mutual information between sensory data and neural representations, with the most common cost in the retinal case being the energetic cost of action potentials transmitted by the RGCs. Despite its simplicity, this principle has proven useful, predicting the center-surround structure of RFs [8], the frequency response profile of contrast sensitivity [9], the structure of retinal mosaics [10, 11], the role of nonlinear rectification [12], different spatiotemporal kernels [13], and inter-mosaic arrangements [14, 15].

While previous studies have largely focused on either spatial or temporal aspects of efficient coding, we optimize an efficient coding model of retinal processing in both space and time to *natural videos* [16]. We systematically varied the number of cells available to the system and found that larger numbers of available cells led to more cell types. Each of these functionally distinct types formed its own mosaic of RFs that tiled space. We show that when and how new cell types emerge and form mosaics is the result of tradeoffs between power constraints and the benefits of specialized encoding that shift as more cells are available to the system. We show that cell types begin by capturing low-frequency temporal information and capture increasingly higher-frequency temporal information over larger spatial RFs as new cell types form. Finally, we investigated the relative arrangement of these mosaics and their dependence on noise. We show that mosaic pairs can be aligned or anti-aligned depending on input and output noise in the system [14]. Together, these results demonstrate for the first time how efficient coding principles can explain, even predict, the formation of cell types, and which types are most informative when channel capacity is limited.

## 2 Model

The model we develop is an extension of [14], a retinal model for efficient coding of natural images, which is based on a mutual information maximization objective proposed in [10]. The retinal model takes *D*-pixel patches of natural images 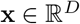 corrupted by input noise 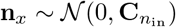, filters these with unit-norm linear kernels 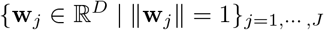 representing *J* RGCs, and then feeds the resulting signals 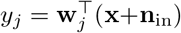 through softplus nonlinearities *η*(*y*) = log (1 + *e^βy^*)/*β* (we used *β* = 0.25) with gain *γ_j_* and threshold *θ_j_*. Finally, these signals are further corrupted by additive output noise 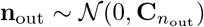, to produce firing rates *r_j_*:

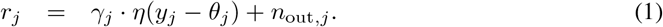

The model learns parameters **w***_j_, γ_j_*, and *θ_j_* to maximize the mutual information between the inputs **x** and the outputs **r**, under a mean firing rate constraint:

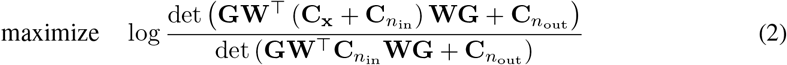

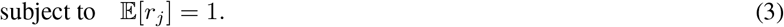

Here **C**_x_ is the covariance matrix of the input distribution, 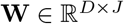 contains the filters **w**_*j*_ as its columns, the gain matrix 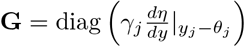, and the noise covariances are 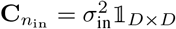 and 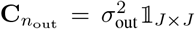. This objective is equivalent to the formulation in [10], which assumes normally distributed inputs and locally linear responses in order to approximate the mutual information in a closed form.

Here, we extend this model to time-varying inputs 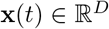 representing natural videos (Figure 1A-B), which are convolved with linear spatiotemporal kernels {**w**_*j*_(*t*)}_*j*=1,···,J_:

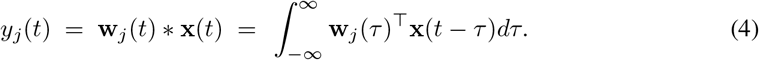

**Figure 1:**
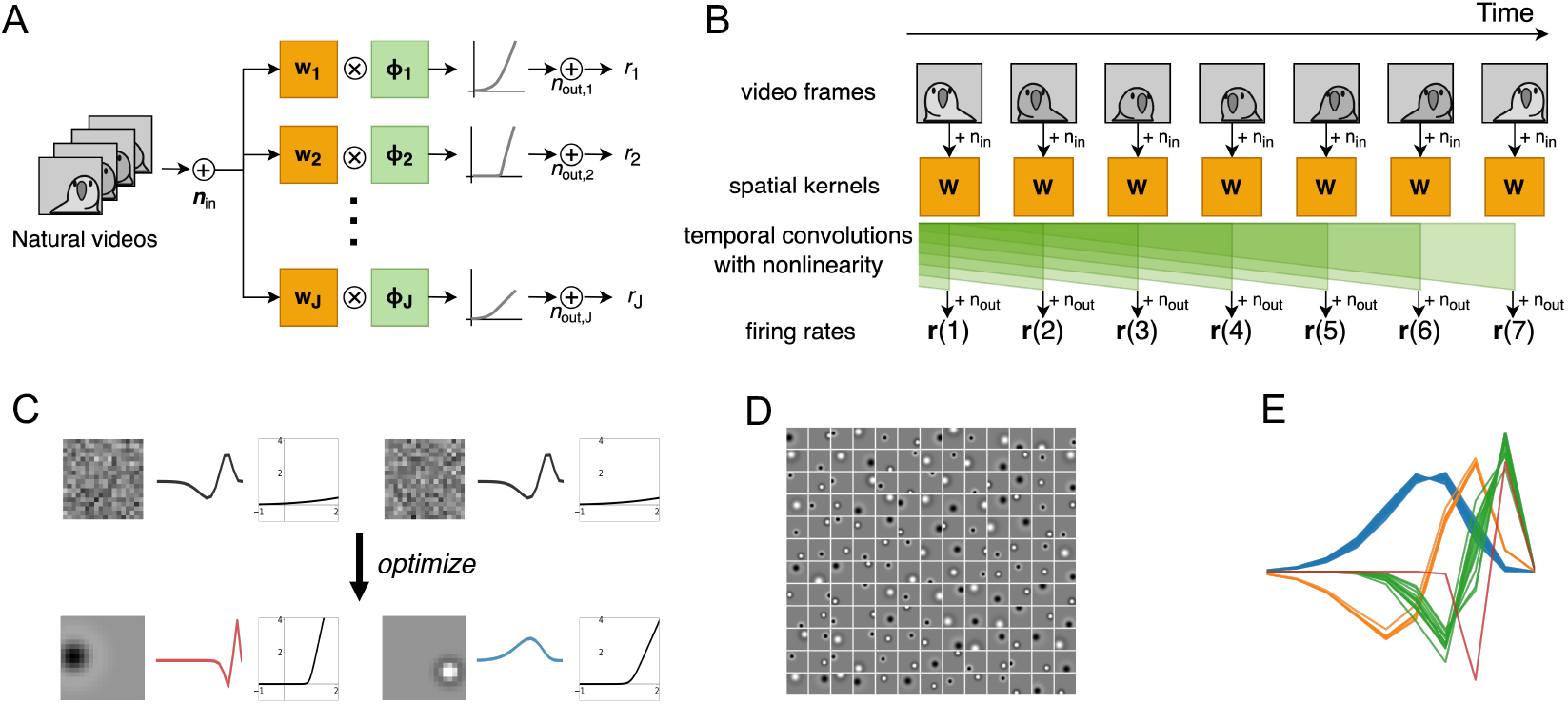
ON and OFF RF mosaics and their temporal RFs are predicted by efficient coding of natural videos. **(A)** Frames of natural videos **x**(*t*) plus input noise **n**_in_ are linearly filtered with the spatial kernels **w**_*j*_ and then passed through one-dimensional temporal convolutions *ϕ_j_* followed by a nonlinearity, resulting firing rates *r_j_*(*t*) for each of *J* RGCs. **(B)** The same calculations shown along the time axis, visualizing the temporal convolutions. **(C)** Examples of initial and optimized spatial filters, temporal filters, and nonlinearities: **(left)** *fast OFF* kernel, **(right)** *slow ON* kernel. **(D)** Unconstrained spatial filters (*J* = 160) learned center-surround shapes, about half of which are ON RFs. **(E)** Temporal filters (*J* = 160) using the parameterization (6) converged to four distinct clusters.

We additionally assume that the convolution kernels are separable in time and space:

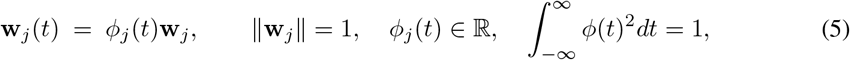

and the temporal kernels are unit-norm impulse responses taking the following parametric form:

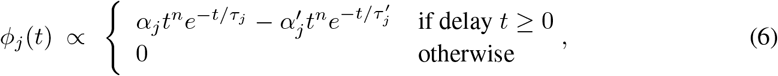

where 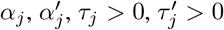 are learnable parameters, and 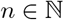 is fixed. Previous work assumed an unconstrained form for these filters, adding zero-padding before and after the model’s image inputs to produce the characteristic shape of the temporal filters in primate midget and parasol cells [13], but this zero-padding represents a biologically implausible constraint, and the results fail to correctly reproduce the observed delay in retinal responses [17, 18, 19]. Rather, optimizing (2) with unconstrained temporal filters produces a filter bank uniformly tiling time (Supplementary Figure 5).

By contrast, (6) is motivated by the arguments of [20], which showed that the optimal minimum-phase temporal filters of retinal bipolar cells, the inputs to the RGCs, take the form

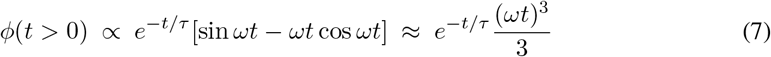

when *ωτ* ≪ 1. Thus, we model RGC temporal filters as a linear combination of these forms. In practice, we take only two filters and use *n* = 6 rather than *n* = 3, since these have been shown to perform well in capturing observed retinal responses [19]. The results produced by more filters or different exponents are qualitatively unchanged (Supplementary Figure 7). For training on video data, we use discrete temporal filters and convolutions with 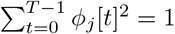. Finally, while unconstrained spatial kernels **w**_j_ converge to characteristic center-surround shapes under optimization of (2) (Figure 1C), for computational efficiency and stability, we parameterized these filters using a radially-symmetric difference of Gaussians

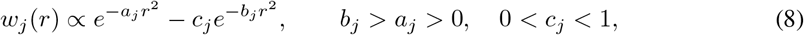

where *r* measures the spatial distance to the center of the RF. The parameters *a_j_, b_j_, c_j_* determining the spatial kernel shape, as well as the center location, are different for each RGC *j*, allowing individual RGCs to have different RF sizes and center-surround strengths. The result of optimizing (2) using these forms is a set of spatial and temporal kernels (Figure 1D-E) that replicate experimentally-observed shapes and spatial RF tiling.

## 3 Efficient coding as a function of channel capacity: linear theory

Before presenting results from our numerical experiments optimizing the model (2, 3), we begin by deriving intuitions about its behavior by studing the case of *linear* filters analytically. That is, We assume a single gain *γ* for all cells, no bias (*θ* = 0), and a linear transfer function *η*(*y*) = *y*. As we will see, this linear analysis correctly predicts the same types of mosaic formation and filling observed in the full nonlinear model. Here, we sketch the main results, deferring full details to Appendix A.

### 3.1 Linear model in the infinite retina limit

For analytical simplicity, we begin by assuming an infinite retina on which RFs form mosaics described by a regular lattice. Under these conditions, we can write the log determinants in (2) as integrals and optimize over the *unnormalized* filter *v* ≡ *γw* subject to a power constraint:

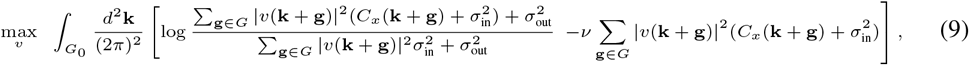

where *C_x_*(**k**) is the Fourier transform of the stationary image covariance *C_x_*(**z** – **z′**), the integral is over all frequencies **k** ∈ *G*_0_ *unique up to aliasing* caused by the spatial regularity of the mosaic, and the sums over **g** account for aliased frequencies (Appendix A.1). In [8], the range [–*π, π*] is used for the integral, corresponding to a one-dimensional lattice and units of mosaic spacing Δ*z* = 1.

Now, solving the optimization in (9) results in a spatial kernel with the spectral form (Appendix A.2)

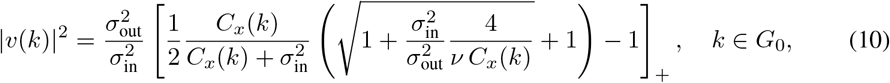

where *k* = ║**k**║ and *ν* is chosen to enforce the constraint on total power. This is exactly the solution found in [8], linking it (in the linear case) to the model of [10, 11]. Note, however, that (10) is only nonzero within *G*_0_, since RF spacing sets an upper limit on the passband of the resulting filters.

The generalization of this formulation to the spacetime case is straightforward. Given a spacetime stationary image spectrum *C*_x_(**z** – **z’**, *t* – *t′*) and radially-symmetric, causal filter *w*(**z**, *t*), the same infinite retina limit as above requires calculating determinants across both neurons *i, j* and time points *t, t′* of matrices with entries of the form

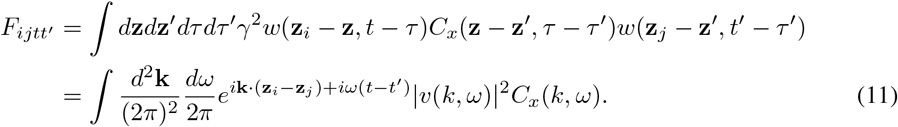

Again, such matrices can be diagonalized in the Fourier basis, with the result that the optimal spacetime filter once again takes the form (10) with the substitutions *v*(*k*) → *v*(*k, ω*), *C_x_*(*k*) → *C_x_*(*k, ω*) (Appendix A.3). Figure 2A depicts the frequency response of this filter in *d* =1 spatial dimensions, with corresponding spatial and temporal sections plotted in Figures 2B-C.

**Figure 2:**
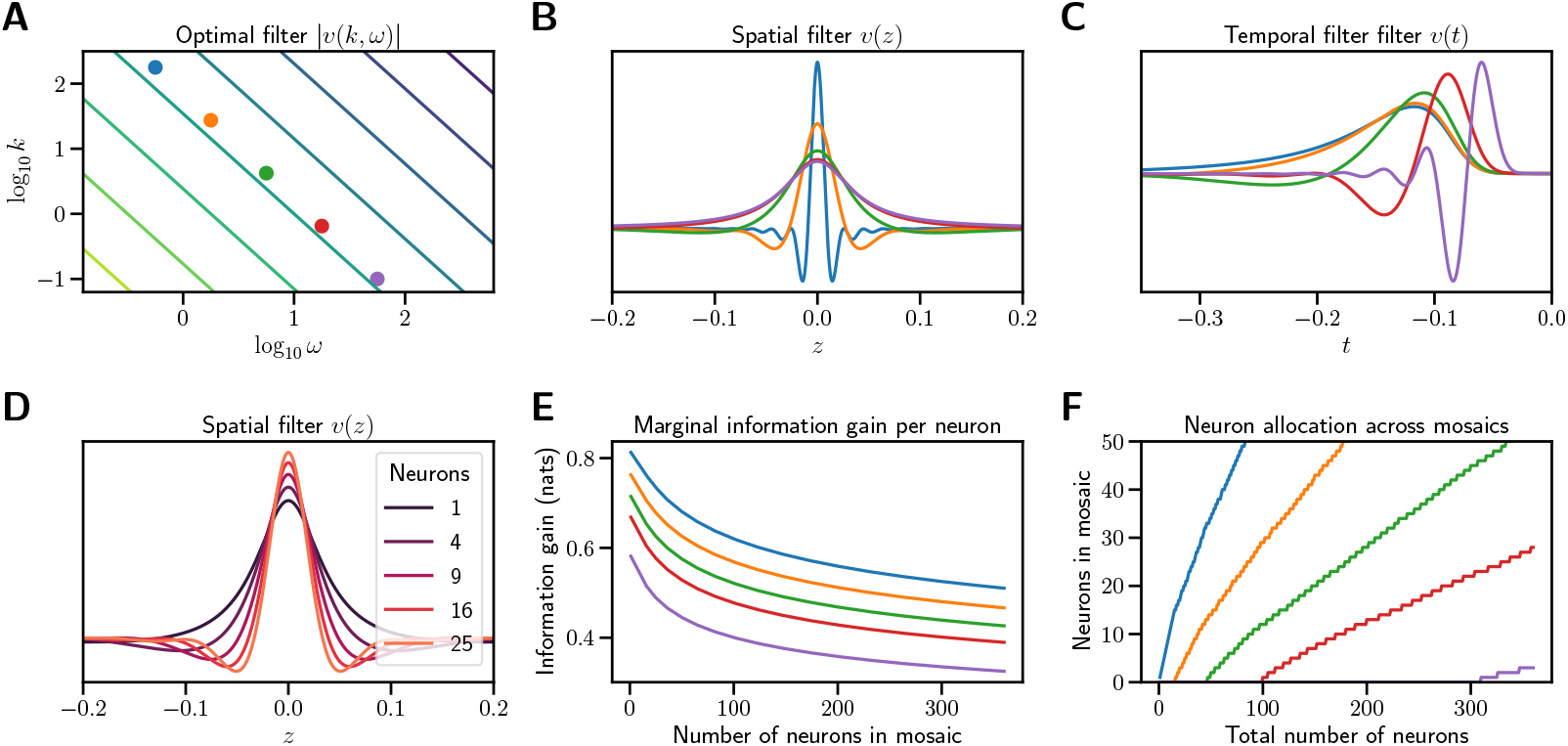
Optimal filters in the linear spacetime case. **(A)** Spectrum of the optimal linear spacetime filter in d =1 spatial dimension. Contour lines indicate constant (log) power. **(B)** Spatial filters at representative temporal frequencies. Each filter represents a vertical section at the correspondingly colored dot in **A**. **(C)** Temporal filters at representative spatial frequencies. Each filter represents a horizontal section at the correspondingly colored dot in **A**. **(D)** Spatial filters at ω = 0 for increasing numbers of RGCs *J*. **(E)** Gain in information per RGC added to each mosaic as a function of current RGC number *J*. New cell types begin when the marginal benefit of adding a RGC to an existing mosaic equals the benefit of adding the first RGC to a new cell type. Color indicates ω_0_, the temporal frequency of the narrow-band filter. **(F)** Total number of RGCs in each mosaic as a function of total RGCs across all mosaics. As new cell types arise and form mosaics, new RGCs are allocated to existing mosaics at a decreasing rate. For both plots, *A* = 100, *σ*_in_ = 0.4, *σ*_out_ = 1.25, and log_10_ *ω*_0_ = 1.5, 1.52, 1.54, 1.56, 1.6. Details of calculations in Appendix A.5.

### 3.2 Multiple cell types and the effects of channel capacity

Up to this point, we have only considered a single type of filter *v*(*k, ω*), corresponding to a single cell type. However, multiple cell types might increase the coding efficiency of the entire retina if they specialize, devoting their limited energy budget to non-overlapping regions of frequency space. Indeed, optimal encoding in the multi-cell-type case selects filters *v* and *v′* that satisfy *v*^*^(*k, ω*)*v′*(*k, ω*) = 0, corresponding encoding *independent* visual information (Appendix A.4).

This result naturally raises two questions: First, how many filter types are optimal? And second, how should a given budget of *J* RGCs be allocated across multiple filter types? As detailed in Appendix A.5, we can proceed by analyzing the case of a finite retina in the Fourier domain, approximating the information encoded by a mosaic of *J* RGCs with spatial filters given by (10) and nonoverlapping bandpass temporal filters that divide the available spectrum (e.g., Figure 2B, C). Following [21], we approximate the correlation spectrum of images by the factorized power law 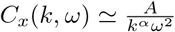 with *α* ≈ 1.3 and find that in this case, the optimal filter response exhibits two regimes as a function of spatial frequency (Supplementary Figure 1A): First, below 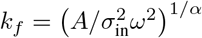, the optimal filter is separable and log-linear, and the filtered image spectrum is white:

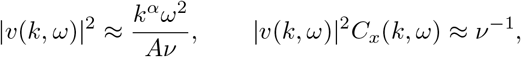

where *ν*, the Lagrange multiplier in (9) that enforces the power constraint, scales as 1/*P* for small values of maximal power *P* and 1/*P*^2^ for larger values (Supplementary Figure 1D). Second, for *k* ≳ *k_f_*, the filter response decreases as *k*^−*α*/2^ until reaching its upper cutoff at 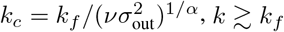, with the filtered image spectrum falling off at the same rate (Supplementary Figure 1B).

But what do these regimes have to do with mosaic formation? The link between the two is given by the fact that, for a finite retina with regularly spaced RFs, adding RGCs decreases the distance between RF centers and so increases the resolving power of the mosaic. That is, the maximal value of k grows roughly as 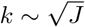 in *d* = 2, such that larger numbers of RGCs capture more information at increasingly higher spatial frequencies (Supplementary Figure 1A). However, while information gain is roughly uniform in the whitening regime, it falls off sharply for *k* ≳ *k_f_* (Supplementary Figure 1C), suggesting the interpretation that the *k* ≲ *k_f_* regime is a “mosaic filling” phase in which information accumulates almost linearly as RFs capture new locations in visual space, while the *k* ≳ *k_f_* regime constitutes a “compression phase” in which information gains are slower as RFs shrink to accommodate higher numbers (Figure 2D). Indeed, one can derive the scaling of total information as a function of *J*:

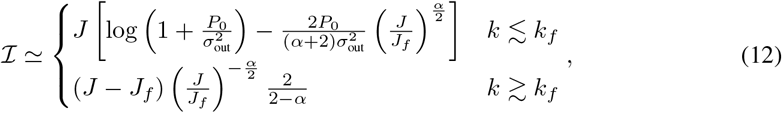

**Figure 3:**
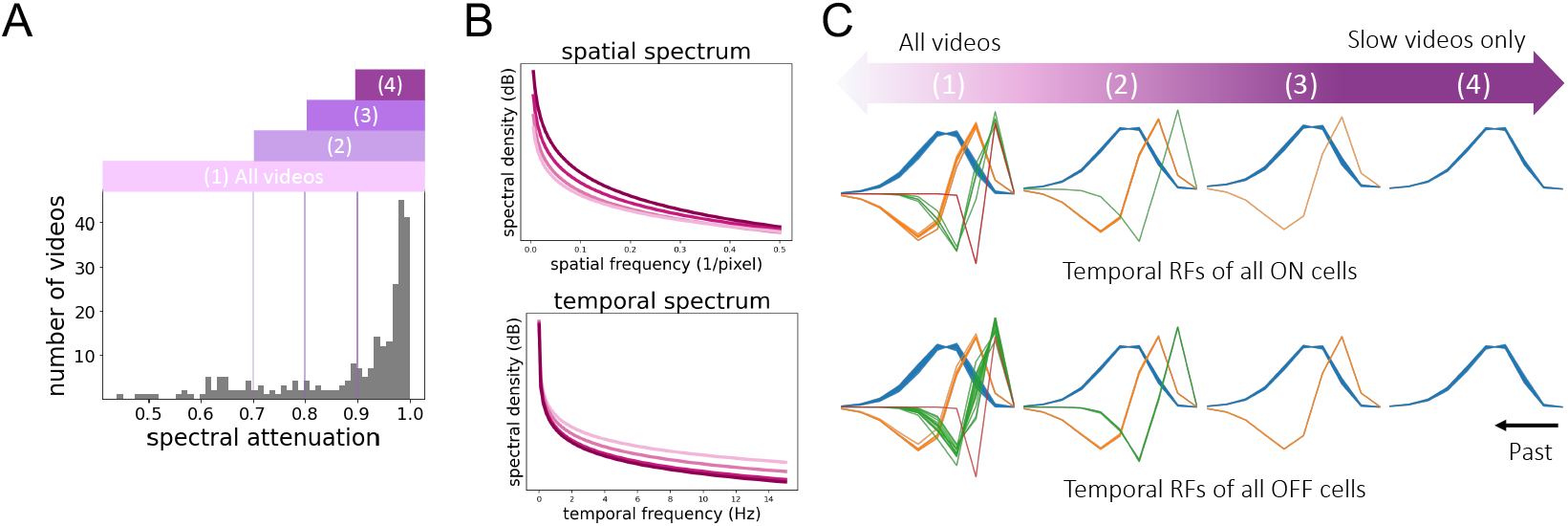
Statistics of natural videos affect learned RFs. **(A)** Histogram of spectral attenuation (fraction of power < 3 Hz) for each video clip from the Chicago Motion Database. A significant portion of the dataset exhibits predominaly low-frequency spectral content in time. Videos with spectral attenuation above 0.9, 0.8, and 0.7, are denoted (4), (3), and (2), respectively, while (1) refers to all videos in the dataset. **(B)** Spatial (top) and temporal (bottom) spectral density of the four subsets. **(C)** Temporal filters learned by training on each of the four subsets. Training on slow videos produced only smoothing kernels, while training on all videos produced a variety of temporal filters.

where *P*_0_ is the power budget per RGC and *J_f_* is the RGC number corresponding to *k* = *k_f_*. Thus, mosaic filling exhibits diminishing marginal returns (Figure 2E), such that new cell types are favored when the marginal gain for growing mosaics with lower temporal frequency drops below the gain from initiating a new cell type specialized for higher temporal frequencies. Moreover, the difference between these gain curves implies that new RFs are not added to all mosaics at equal rates, but in proportion to their marginal information (Figure 2F). As we demonstrate in the next section, these features of cell type and mosaic formation continue to hold in the full nonlinear model in simulation.

## 4 Experiments

We analyzed the characteristics of the optimal spatiotemporal RFs obtained from the model (2, 3) trained on videos from the Chicago Motion Database [22]. Model parameters for spatial kernels, temporal kernels, and the nonlinearities were jointly optimized using Adam [23] to maximize (2) subject to the mean firing rate constraint (3) using the augmented Lagrangian method with the quadratic penalty *ρ* = 1 [24]. Further technical details of model training are in Appendix E.

As previously noted, the power spectral density of natural videos can be well approximated by a product of spatial and temporal power-law densities, implying an anticorrelation between high spatial and temporal frequency content [21]. Supplementary Figure 4 shows the data spectrum of the videos in our experiments is also well-approximated by separable power-law fits. To examine the effect of these statistics on the learned RFs, we divided the dataset into four progressively smaller subsets by the proportion of their temporal spectral content below 3 Hz, their *spectral attenuation*. Using values of 70%, 80%, and 90% then yielded a progression of datasets ranging from most videos to only the slowest videos (Supplementary Figure 3A, B). Indeed, when the model was trained on these progressively slower data subsets, it produced only temporal smoothing filters, whereas the same model trained on all videos produced a variety of “fast” temporal filter types (Supplementary Figure 3C). We also note that these experiments used *unconstrained* spatial kernels in place of (8), yet still converged on spatial RFs with typical center-surround structure as in [10, 15, 14]. Thus, these preliminary experiments suggest that the optimal encoding strategy—in particular, the number of distinct cell types found—depends critically on the statistics of the video distribution to be encoded.

### 4.1 Mosaics fill in order of temporal frequency

As the number of RGCs available to the model increased, we observed the formation of new cell types with new spectral properties (Figure 4). We characterized the learned filters for each RGC in terms of their spectral centroid, defined as the center of mass of the Fourier (spatial) or Discrete Cosine (temporal) transform. Despite the fact that each model RGC was given its own spatial and temporal filter parameters (8, 6), the learned filter shapes strongly clustered, forming mosaics with nearly uniform response properties (Figure 4A–C). Critically, the emergence of new cell types shifted the spectral responses of previously established ones, with new cell types compressing the spectral windows of one another as they further specialized. Moreover, mosaic density increased with increasing RGC number, shifting the centroids of early mosaics toward increasingly higher spatial frequencies. This is also apparent in the forms of the typical learned filters and their power spectra: new filters selected for increasingly high-frequency content in the temporal domain (Figure 4D).

**Figure 4:**
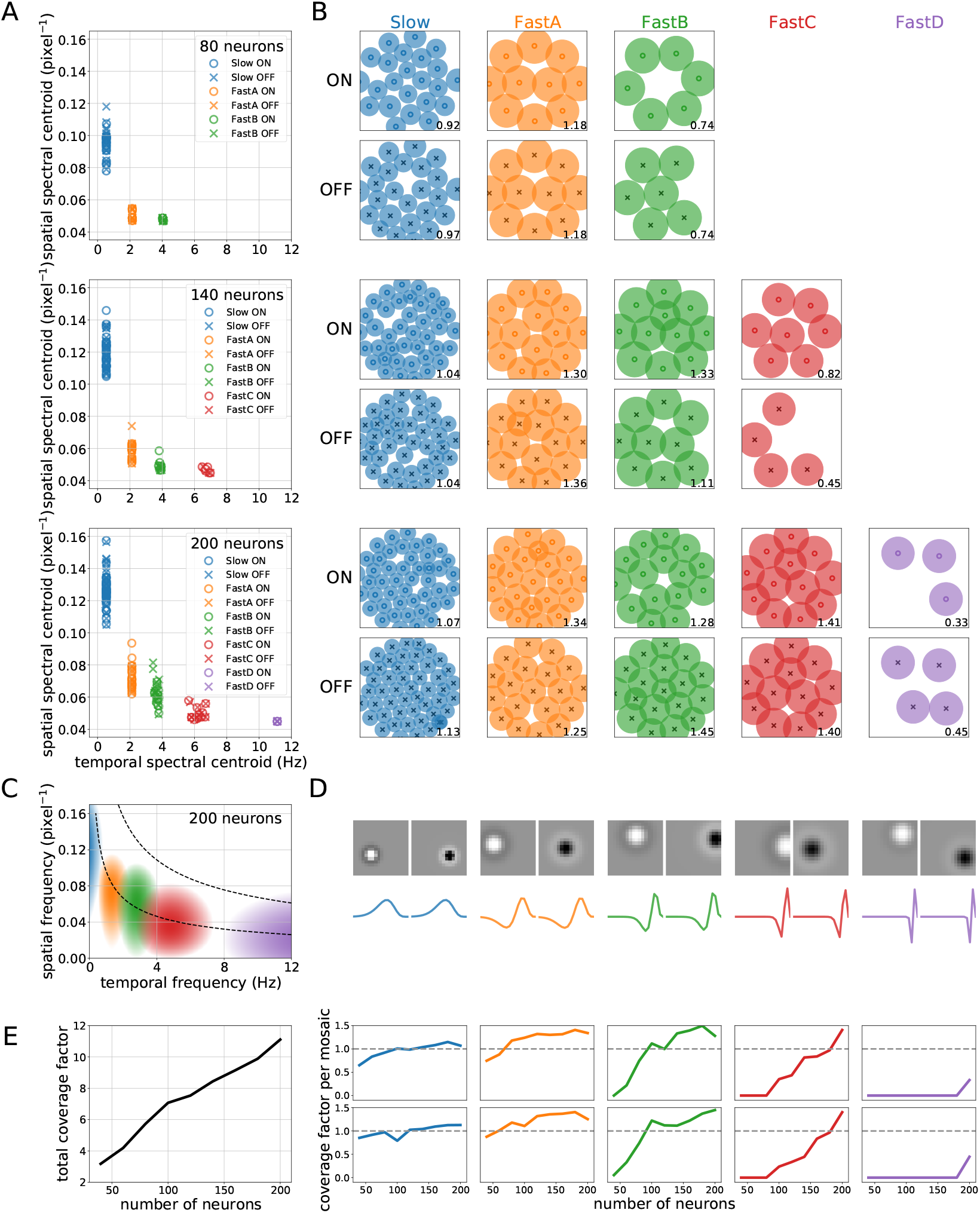
Emergence of new RF types with increasing RGC number. **(A)** Distribution of spatial and temporal spectral centroids for *J* = 80, 140, 200 RGCs. ON and OFF RFs form distinct clusters corresponding to different learned filter types. **(B)** ON and OFF mosaics corresponding to each cell type. The number in the lower right of each plot is the coverage factor for the mosaic. **(C)** Power spectral density of a typical kernel in each mosaic for *J* = 200. As predicted, learned kernels filter over roughly nonoverlapping regions of spatiotemporal frequency. Contour lines represent isopower lines of the signal correlation *C_x_*(*k, ω*). **(D)** Learned shapes of example spatial ON and OFF filters (top) and corresponding temporal filters (bottom) from each RF type for the *J* = 200 case. **(E)** Total (left) and per-mosaic (right) coverage factors as the number of RGCs *J* increases from 40 to 200. New mosaics increase coverage linearly with the number of RFs, while nearly full mosaics see diminishing returns in coverage from density increases. See Supplementary Figures 8–9 for similar plots for all RGC numbers.

We likewise analyzed the coverage factors of both individual mosaics and the entire collection, defined as the proportion of visual space covered by the learned RFs. More specifically, we defined the spatial radius of an RF as the distance from its center at which intensity dropped to 20% of its peak and used this area to compute a coverage factor, the ratio of total RF area to total visual space (*π*/4 of the square’s area due to circular masking). Since coverage factors depend not simply on RGC number but on RF density, they provide an alternative measure of the effective number of distinct cell types learned by the model. As Figure 4E shows, coverage increases nearly linearly with RGC number, while coverage for newly formed mosaics increases linearly before leveling off. In other words, new cell types initially increase coverage of visual space by adding new RFs, but marginal gains in coverage diminish as density increases. In all cases, the model dynamically adjusts the number of learned cell types and the proportion of RGCs assigned to them as channel capacity increases.

### 4.2 Phase changes in mosaic arrangement

In addition to retinal organization at the level of mosaics, a pair of recent papers reported both experimental [15] and theoretical [14] evidence for an additional degree of freedom in optimizing information encoding: the relative arrangement of ON and OFF mosaics. Jun et al. studied this for the case of natural images in [14], demonstrating that the optimal configuration of ON and OFF mosaics is alignment (RFs co-located) at low output noise levels and anti-alignment (OFF RFs between ON RFs and vice-versa) under higher levels of retinal output noise. Moreover, this transition is abrupt, constituting phase change in optimal mosaic arrangement.

We thus asked whether learned mosaics exhibited a similar phase transition for natural video encoding. To do so, following [14], we repeatedly optimized a small model (*J* = 14, 7 ON, 7 OFF) for multiple learned filter types while systematically varying levels of input and output noise. In each case, one ON-OFF pair was fixed at the center of the space, while the locations of the others were allowed to vary. We used RF size *D* = 8^2^ pixels for *Slow* and *D* = 12^2^ for *FastA* and *FastB* to allow the size of spatial kernels to be similar to those of the previous experiments, and we imposed the additional constraint that the shape parameters *a_j_, b_j_*, and *c_j_* in (8) are shared across RGCs.

Under these conditions, the six free pairs of RFs converged to either aligned (overlapping) or anti-aligned (alternating) positions along the edges of the circular visual space, allowing for a straightforward examination of the effect of input and output noises on mosaic arrangement. Figure 5A-C shows that the phase transition boundaries closely follow the pattern observed in [14]: increasing output noise shifts the optimal configuration from alignment to anti-alignment. Moreover, for each of the tested filters, increasing input noise *discourages* this transition. This effect also follows from the analysis presented in [14], since higher input noise increases coactivation of nearby pairs of RFs, requiring larger thresholds to render ON-OFF pairs approximately indpendent (Appendix B).

**Figure 5:**
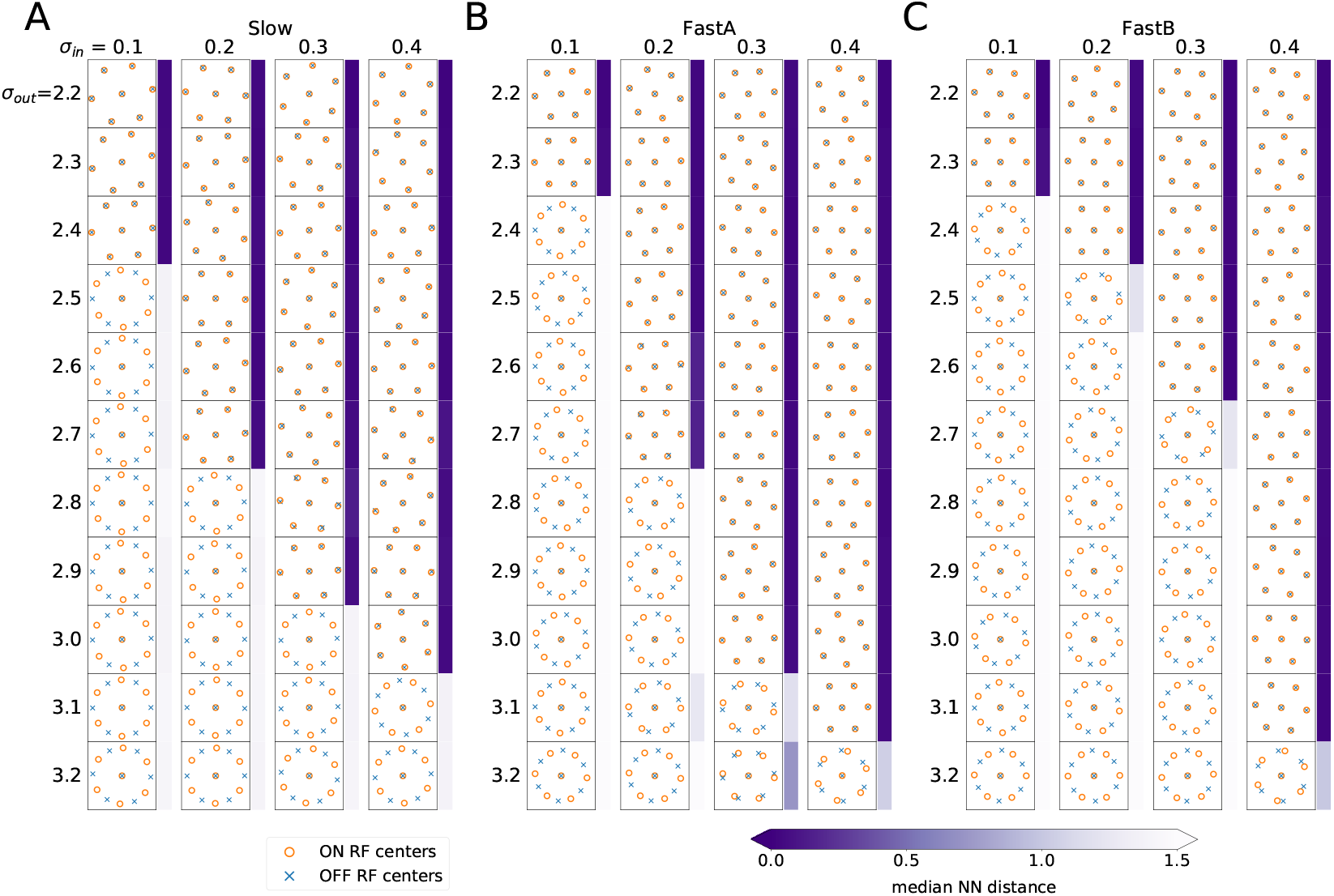
Learned mosaics exhibit a phase transition as a function of input and output noise. **(A-C)** Spatial kernel centers for *Slow* **(A)**, *FastA* **(B)**, and *FastB* **(C)** as a function of *σ*_in_ and *σ*_out_. In all three cases, the optimal configuration changes from aligned to anti-aligned when output noise increases or input noise decreases. Blue bars denote alignment as measured by median distance to the nearest RF center of the opposite type.

## 5 Discussion

### Related work

As reviewed in the introduction, this study builds on a long line of work using efficient coding principles to understand retinal processing. In addition, it is related to work examining encoding of natural videos [25, 22, 16] and prediction in space-time. The most closely related work to this one is that of [13], which also considered efficient coding of natural videos and considered the tradeoffs involved in multiple cell types. Our treatment here differs from that work in several key ways: First, while [13] was concerned with demonstrating that multiple cell types could prove beneficial for encoding (in a framework focused on reconstruction error), that study predetermined the number of cell types and mosaic structure and only optimized their relative spacing. By contrast, this work is focused on how the number of cell types is dynamically determined, and how the resulting mosaics arrange themselves, as a function of the number of units available for encoding (i.e., the channel capacity). Specifically, we follow previous efficient coding models [8, 9, 10, 11] in maximizing mutual information and do not assume an a priori mosaic arrangement, a particular cell spacing, or a particular number of cell types—all *of these emerge via optimization in our formulation*. Second, while the computational model of [13] optimized strides for a pair of rectangular arrays of RGCs, we individually optimize RF locations and shapes, allowing us to study changes in optimal RF size and density as new, partial mosaics begin to form. Third, while [13] used zero-padding of natural videos to bias learned temporal filters toward those of observed RGCs, we link the form of temporal RFs to biophysical limits on the filtering properties of bipolar cells, producing temporal filters with the delay properties observed in real data. Finally, while [13] only considered a single noise source in their model, we consider noise in both photoreceptor responses (input noise) and RGC responses (output noise), allowing us to investigate transitions in the optimal relative arrangement of mosaics [14, 15].

We have shown that efficient coding of natural videos produces multiple cell types with complementary RF properties. In addition, we have shown for the first time that the number and characteristics of these cell types depend crucially on the channel capacity: the number of available RGCs. As new simulated RGCs become available, they are initially concentrated into mosaics with more densely packed RFs, improving the spatial frequency bandwidth over which information is encoded. However, as this strategy produces diminishing returns, new cell types encoding higher-frequency temporal features emerge in the optimization process. These new cell types capture information over distinct spatiotemporal frequency bands, and their formation leads to upward shifts in the spatial frequency responses of previously formed cell types. Moreover, pairs of ON and OFF mosaics continue to exhibit the phase transition between alignment and anti-alignment revealed in a purely spatial optimization of efficient coding [14], suggesting that mosaic coordination is a general strategy for increasing coding efficiency. Furthermore, despite the assumptions of this model—linear filtering, separable filters, firing rates instead of spikes—our results are consistent with observed retinal data. For example, RGCs with small spatial RFs exhibit more prolonged temporal integration: they are also more low-pass in their temporal frequency tuning. Second, there is greater variability in the size and shape of spatial RFs at a given retinal location, but temporal RFs exhibit remarkably little variability in our simulations and in data [19]. Thus, these results further testify to the power of efficient coding principles in providing a conceptual framework for understanding the nervous system.

## Acknowledgments and Disclosure of Funding

This work was supported by NIH/National Eye Institute Grant R01 EY031396.

## A Analysis details for the linear model

### A.1 Derivation of the objective in the infinite retina limit

Here, we provide details for the derivation of (9). To begin, we assume that spatial RFs centered at locations **z**_*i*_, are circularly symmetric: *w*(**z**_*i*_, **z**) = *w*(|**z**_i_ – **z**|). Moreover, we assume that the RF centers form a Bravais lattice *R* with basis vectors (**a**_1_, **a**_2_) in *d* = 2 such that **z**_*i*_ = *n*_1*i*_**a**_1_ + *n*_2*i*_**a**_2_ for some integers (*n*_1*i*_, *n*_2*i*_) and the mosaic is translationally invariant under these shifts. Likewise, there exist basis vectors (**b**_1_, **b**_2_) for the dual lattice *G* such that **a**_*i*_ · **b**_*j*_ = 2*πδ_ij_*, with the interpretation that while the RF mosaic is invariant under shifts by integer multiples of **a**_*i*_, its representation in Fourier space is invariant under shifts by the **b**_*i*_. Note that this is distinct from the assumption that the encoded images themselves are invariant under such translations. Here, for simplicity, we will transition to an infinite retina, though we will relax this assumption later.

Now, consider the determinants in (2). In the linear, discrete purely spatial case, the numerator contains a determinant over *J* × *J* matrices with elements

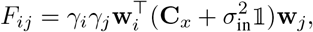

which can be written as

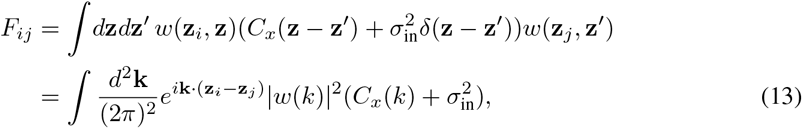

in the continuous case, where we have made use of the translational and rotational symmetry of both *w* and *C_x_* to write their Fourier transforms in terms of *k* = ||**k**||. Of course, in the continuum case, *J* → *∞*, and we have for a symmetric, positive-definite matrix **A**

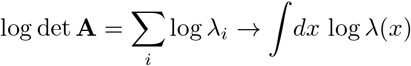

with λ the eigenspectrum of **A** in the continuum limit.

Fortunately, the continuum expression (13) can be diagonalized in the Fourier basis. Let *ψ_j_* (**k**′) = *e*^i**k**′·**z**_*j*_^. Then

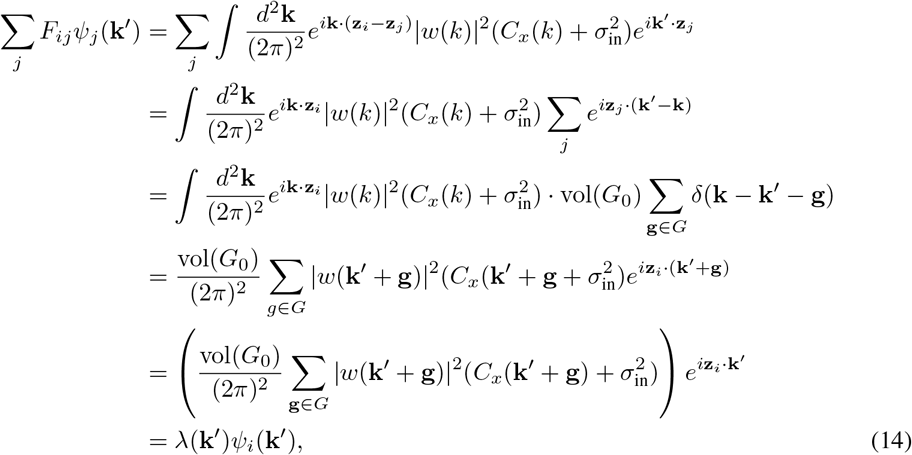

where again 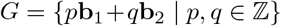 is the dual lattice, 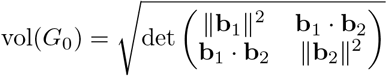 is the volume of the unit cell of the dual lattice, and **z**_*i*_ · **g** is an integer multiple of 2*π* by definition of the lattices *R* and *G*. Likewise, we note that, due to aliasing from the mosaic spacing, *ψ*(**k** + **g**) = *ψ*(**k**), so the only unique eigenvalues are those with **k** ∈ *G*_0_, the unit cell of the dual lattice. Thus, by a similar calculation for the denominator, the terms in (2) become

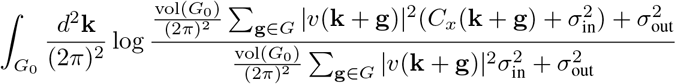

in the continuum limit.

As for the constraint term, the restriction (3) fails to generalize to the linear case, where 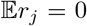. Instead, we note that for (1), we have

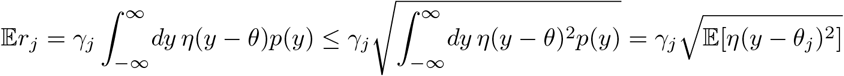

by Hölder’s inequality. That is, we can enforce a looser restriction on firing rates by bounding the power used by the filter. Of course, in the linear case considered above, *θ* = 0, *η*(*y*) = *y*, and 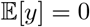, so that the inequality becomes

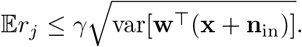

In practice, we bound the square of this expression, which yields the continuous objective (9).

### A.2 Derivation of the optimal linear filter

As noted in Section 3.1 the optimal solution for (9) takes the form (10). However, this expression differs in two key aspects from the form originally presented in [8]: First, (10) involves a rectification operation [·]_+_ on the coefficients of the filter. Second, the integral over **k** is over the unit cell of the dual lattice *G*_0_, not over the interval [−*π*, *π*] (in dimension *d* = 1). Here, we provide details pertaining to both of these points.

Our starting point is (9). Here, to simplify notation, we work in dual lattice units such that vol(*G*_0_) = (2*π*)^2^, since we can restore this later by the transformation 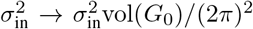, *C_x_* → *C_x_*vol(*G*_0_)/(2*π*)^2^. Moreover, as in [8], we recognize that the optimization objective is a function only of the power spectrum |*ν*(**k** + **g**) |^2^. Varying with respect to this quantity, however, requires that we enforce a positivity constraint, which implies a modified objective

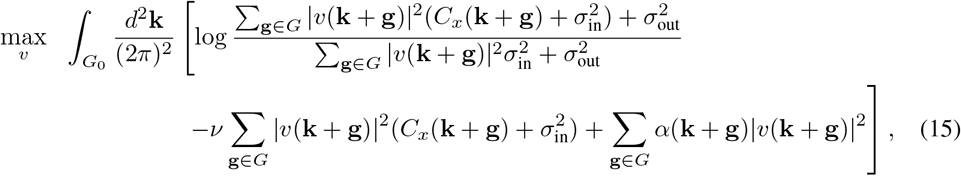

where *α*(**k** + **g**) is a Lagrange multiplier (one per frequency) enforcing the positive-definiteness of the filter power. Taking derivatives with respect to |*ν*(**k** + **g**) |^2^ and rearranging then gives:

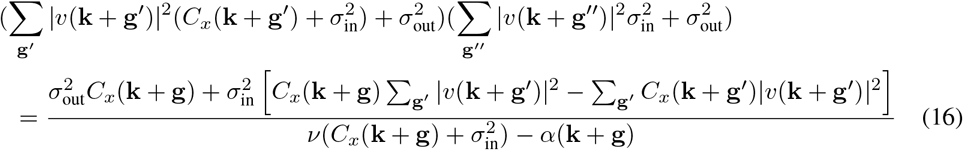

supplemented by the complementary slackness conditions *α*(**k** + **g**)|v(**k** + **g**)|^2^ = 0.

Two things are important to note about this equation: First, if the sums over **g** are reduced to single terms (i.e., there is no power in either the correlation or filter spectra outside the unit cell of the reciprocal lattice), the term in brackets in the numerator of the right-hand side vanishes. If, in addition, *α* = 0, we are back with the same expression as found in [8]. Second, the left-hand side of this equation is *the same for all* **g** ∈ *G*, which implies that the right-hand side must be as well. Thus,

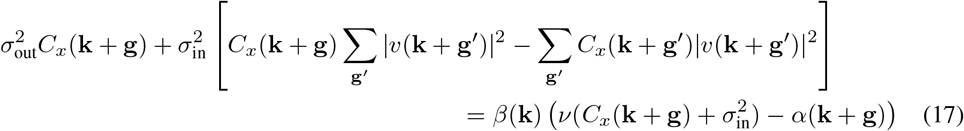

with *β*(**k**) > 0. Recall here that, with respect to the optimization, *C_x_*, 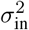, and 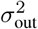 are constants and *β*(**k**) is a function of |*ν*(**k** + **g**) |^2^ and these constants. Only |*ν*(**k** + **g**) |^2^ and α(**k** + **g**) are variables (and these are tied by complementary slackness). This suggests that the term in brackets cannot vanish for general *C_x_* unless only a single *ν*(**k** + **g**′) in the sum is nonzero.

We check this solution in three cases:

1. Assume *ν*(**k** + **g**) ≠ 0 and *ν*(**k** + **g**′) = 0 for **g**′ ≠ **g**. Then the term in brackets vanishes, complementary slackness dictates *α*(**k** + **g**) = 0, and the situation reduces to (19) below.
2. Assume *ν*(**k** + **g**) = 0 but *ν*(**k** + **g**′) ≠0 for a single **g**′ ≠ **g**. Then complementary slackness allows *α* (**k** + **g**) ≠ 0 and we have

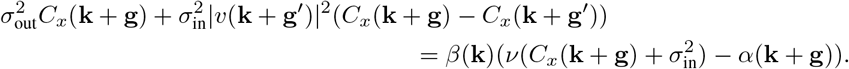 Of course, if we evaluate the right-hand side of (16) at **g** = **g**′, we get (assuming *ν* > 0)

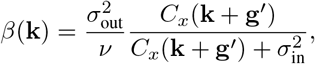

and we can rewrite the optimality condition as

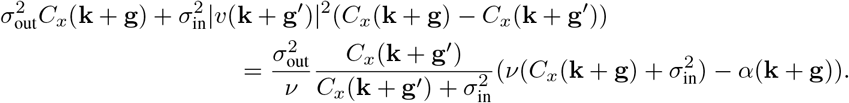 If we divide both sides by 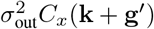, we see that a solution with *α*(**k** + **g**) ≥ 0 always exists provided

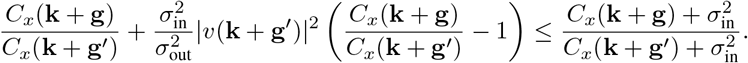 Now when 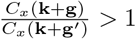, the middle term in the inequality is positive but the first term on the left is larger than the right-hand side, so the inequality can never hold. By contrast, when 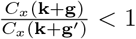, the middle term is negative and the first term on the left is already smaller than the right-hand side, and the inequality always holds. Thus, the condition we need is that

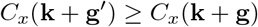

for all **g** ∈ *G*. That is, the filter should respond only at the (aliased) frequency with highest power.
3. Assume *ν*(**k** + **g**) = 0 for all *g* ∈ *G*. This happens, for instance, when the RF filter has a maximum frequency response at *k*_max_, which gives *ν*(**k** + **g**) =0 for all ||**k**|| > *k*_max_. Then, from (16) first-order stationarity requires

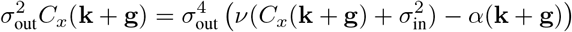

or

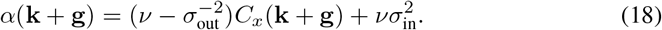

Together, these calculations suggest the following pattern in the filter response *ν*(**k** + **g**): Assuming that *C_x_*(*k*) is a monotonically decreasing function of *k* = ||*k*||, the filter can only respond for at most *one* aliased value of the frequency **k**. The optimal frequency at which to respond is, from point 2 above, the one with the highest power, and given our assumption about *C_x_(**k***), this is the one with the *lowest frequency*. As a result, *ν*(**k**) only has support within the unit cell *G*_0_, the sums over **g** ∈ *G* vanish, and at these frequencies, the solution is given by the term inside the brackets in (10). Moreover, at values of **k** for which the term inside the brackets in (10) would be 0 or negative, the correct solution is given by *ν*(**k** + **g**) = 0 with α(**k** + **g**) given by (18). Thus, returning to (16), we have

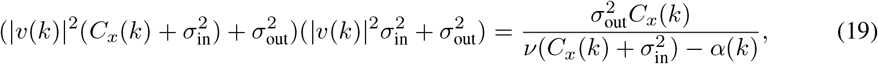

and collecting terms gives

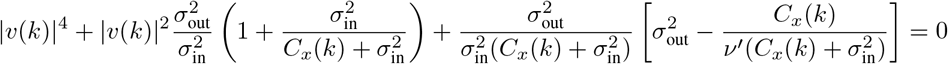

with 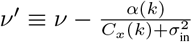. This has the solution

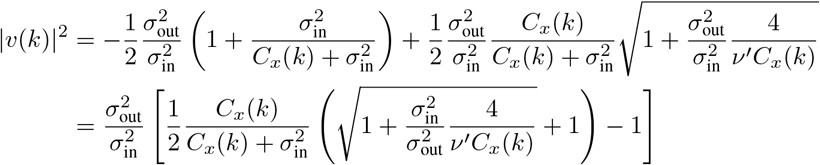

When the term in brackets is positive, complementary slackness requires *α*(*k*) = 0 and thus *ν*′ = *ν*, while when the bracketed term is less than or equal to zero, *α*(*k*) need not be zero, but complementary slackness requires |*ν*(*k*) |^2^ = 0, which holds when *α*(*k*) satisfies (18). Thus, the full solution is given by (10).

### A.3 Extenstion to the spatiotemporal case

Consider the same setup as in the previous section but with a spacetime filter *γw*(**z**, *t*) = *ν*(**z**, *t*). Now, the relevant correlations inside the determinant, analogous to (13), are between firing rates at different filters at different times:

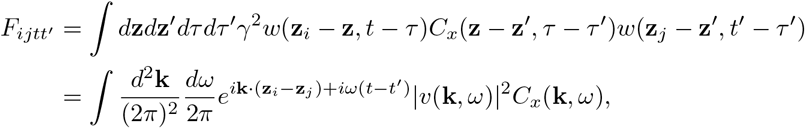

and once again we can diagonalize this by taking eigenvectors

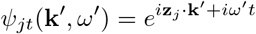

for which

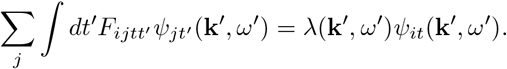

Now, just as in the spatial domain, this leads to an objective in the Fourier domain

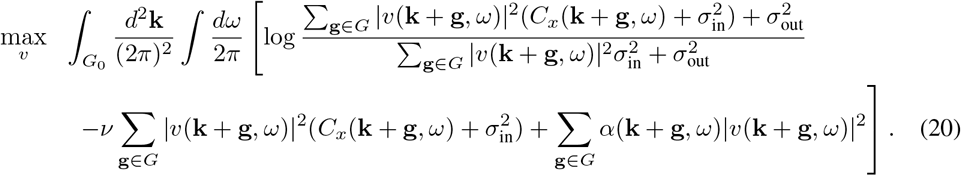

Clearly, by the same arguments as above, the solution to this is once again (10) with *ν*(*k*) → *ν*(*k,ω*), subject to a normalization condition over both spatial and temporal frequencies.

However, the solution given in (10) only determines the power spectrum of the optimal filter, not its phase. That is, if *ν*(**k**, *ω*) = |*ν*(**k**, *ω*)|*e*^*iϕ*(**k**, *ω*)^, *ϕ*(**k**, *ω*) is undetermined. However, for the minimum-phase system, which is both causal and imposes the minimum temporal delay between the incoming signal and its filtered response [20], there is a relation between the filter’s amplitude |*ν*(**k**, *ω*) | (treating **k** as constant) and its phase *ϕ*(**k**, *ω*):

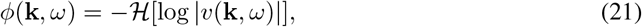

where 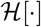 is the Hilbert transform (over *ω*) [26]. More explicitly, the minimum phase filter has

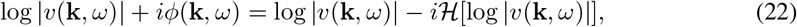

which is the complex conjugate of the analytic signal and so is analytic in the *lower* half plane (and thus causal). For a given, fixed **k**, this can be used to plot the causal temporal filters as depicted in Figure 2C. Note also that the spatiotemporal filter found by generalizing (10) is not exactly separable, though it may be well approximated by the product of a spatial and temporal filter in certain regimes.

Finally, we note that the solution (19) in both its spatial and spatiotemporal forms exhibits a kink in its power spectral density stemming from the positive rectification. This in turn implies that the optimal spatial filters exhibit ringing in both the space and time domains, unlike the actual observed retinal filters. Of course, these solutions, like many ideal filters, are all but impossible to implement in real systems, necessitating tradeoffs among ripple, rolloff, and other attributes [26]. Indeed, as noted above, allowable temporal filters are constrained by bipolar cell response properties [20]. As a result, in Figure 2B-D, we have smoothed the optimal power spectral densities with a double exponential kernel *e*^−*a*|*k*|^ (*e*^−*α*|*ω*|^) prior to transforming back to the space (time) domain.

### A.4 Independence of multiple filters

Here, we generalize the approach given in Appendix A.1 to multiple filter types, arguing that requiring these filters not overlap in frequency space is sufficient to optimize the objective 2 in the infinite retina limit. The only relevant difference between this case and that of Appendix A.1 is that, in addition to terms like (13), the determinants also contain cross-terms of the form

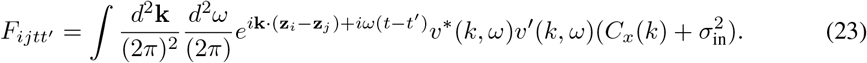

The presence of such cross-terms, which arises from correlations between activity in the two mosaics, reduce the magnitude of the determinants and thus overall information. However, we argue that such cross-terms should vanish for the optimal solution, yielding an information objective that is equivalent to a sum of individual mosaic terms like (9). Intuitively, this is because our power constraint in (9) remains unchanged under an orthogonal transformation of the filters, while information is *subadditive* when cross-terms between filters are nonzero. Thus, we can always increase information by choosing a filter basis in which both **C**_*x*_ and **C**_in_ are block diagonal and the mosaics decouple. This logic is similar to the argument of [13], where it was reconstruction performance that was equivalent under transformations of the filters but costs were superadditive.

However, this reasoning does not fully fix the choice of independent filters, since there are multiple ways to make off-diagonal blocks in the determinants vanish. For example, in the discrete (pixel, frame) formulation of the problem, the generalized eigenvectors of (**C**_*x*_, **C**_in_) represent a special solution (the one that *fully* diagonalizes both matrices). Of course, these filters need not optimize the information objective. But there is an alternative solution that removes only the off-diagonal blocks in the determinants, reducing the problem to one of again optimizing individual filters with the objective (9): choosing spectrally disjoint filters with *ν*\* (*k, ω*)*ν*′(*k, ω*) = 0 for different mosaics defined by *ν* and *ν*′. Note that this condition is merely sufficient, not necessary, to decouple mosaics, but it is independent of any assumptions on the image correlation structure **C**_*x*_ or the filters *ν*(*k, ω*). More specifically, it does not depend on any assumptions of spacetime separability for either.

In [13], the authors considered a form of this same bandwidth partitioning in the spatial case (*ν*\* (*k*)*ν*′(*k*) = 0), but as we show Appendix A.5, this approach encounters a limit beyond the first two mosaics, when the spatial passband spectra of new filters substantially overlap. Rather, the more general solution is that mosaic filters are band-limited in both space and time, arranging themselves to tile minimally overlapping regions with highest power in the (*k, ω*) plane, as we find in the full nonlinear model (Figure 4).

### A.5 Effects of channel capacity and new mosaic formation

Now we reconsider the analysis of Appendices A.1 and A.3 in the case that the number of RGCs *J* in the system is varied. To do so, we take a *finite* retina of size (*L*_1_, *L*_2_) = (*M*_1_ ║**a**_1_║, *M*_2_ ║**a**_2_║) along each basis vector of the lattice R and assume a periodic extension of the signal outside this domain. This approximation allows us to continue working in the Fourier domain but sets a *lower* bound on the spatial frequencies a mosaic can hope to resolve (in addition to the *upper* bound resulting from the mosaic lattice spacing). Note that for a fixed retina size, increasing *M_i_* is equivalent to reducing ║**a**_*i*_║, i.e., decreasing the spacing between RF centers.

More explicitly, let **Z**_*j*_ = *n*_1*j*_**a**_1_ + *n*_2*j***a**_2__ as before, with RGC number *J* ≈ vol(*R*)/vol(*R*_0_) = *M*_1_*M*_2_. That is, the number of RGCs is the size of the retina divided by the unit cell volume of the Bravais lattice. In addition, the assumed periodicity of the signals implies that Fourier transforms involve only frequencies of the form 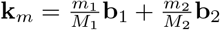 for 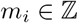. From this, following the derivation preceding (14), we have that *ψ_j_* (**k**_*l*_) = *e*^*i***Z**_*j*_·*k*_*l*_^ are eigenvectors of the *J* × *J* matrix *F_ij_*:

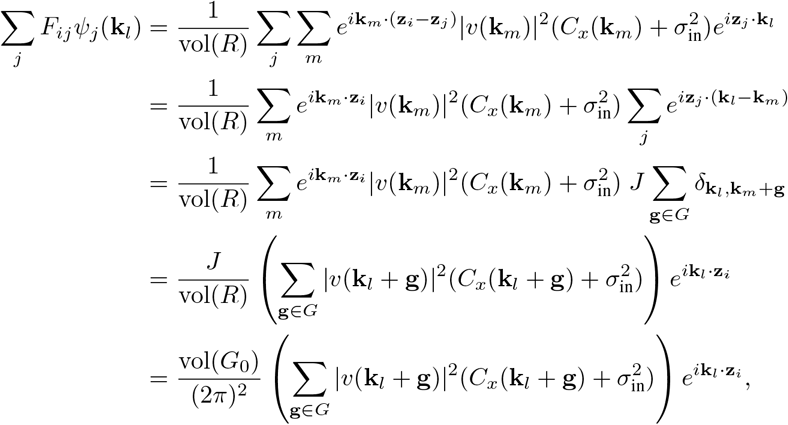

since vol(*R*_0_) · vol(*G*_0_) = (2*π*)^2^ ⟹ *J*/vol(*R*) = 1/vol(*R*_0_) = vol(*G*_0_)/(2*π*)^2^. Clearly, this is the analogue of (14), and the form of the optimal spatial filter 10 once again holds in spacetime, albeit only at a finite set of frequencies **k**_*m*_ ∈ *G*_0_. In what follows, we will be interested in the effect of changing J on the information encoded by the filters.

Turning to the images themselves, we here consider an idealized form of the power spectrum of natural videos [21],

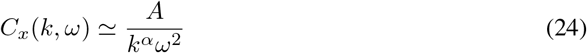

where *α* ≈ 1.3. Of course, this is not the same as the spectrum of images presented to the retina, since the latter is low-pass filtered via the modulation transfer function of the eye [9], but here we analyze this simpler form, since the qualitative results are similar in both cases. For this spectrum, the optimal filter (10) takes the form

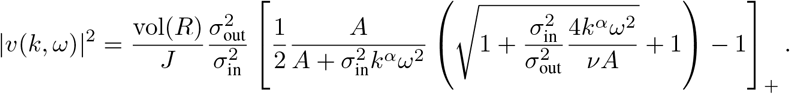

Now, we will define two useful frequencies in terms of our parameters

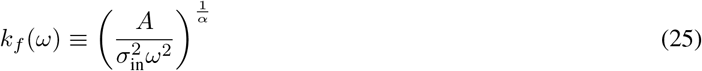

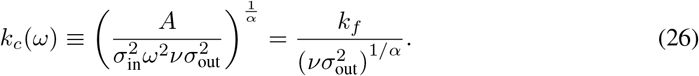

Often, 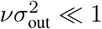, so that *k_c_* ≫ *k_f_*. In terms of these, the filter can be rewritten

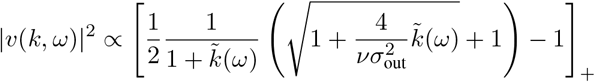

with 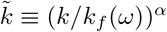. That is, up to an overall rescaling, the filter depends only on the ratio 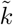 and the dimensionless parameter 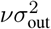. Naively, changes in *k_f_* simply rescale the filter in space, while *ν* is related to the power consumed by the filter. However, given a *fixed* power budget, these two parameters are tied, leading to a single family of spatial filters that depend on the temporal frequency *ω* (and vice-versa).

**Supplementary Figure 1:**
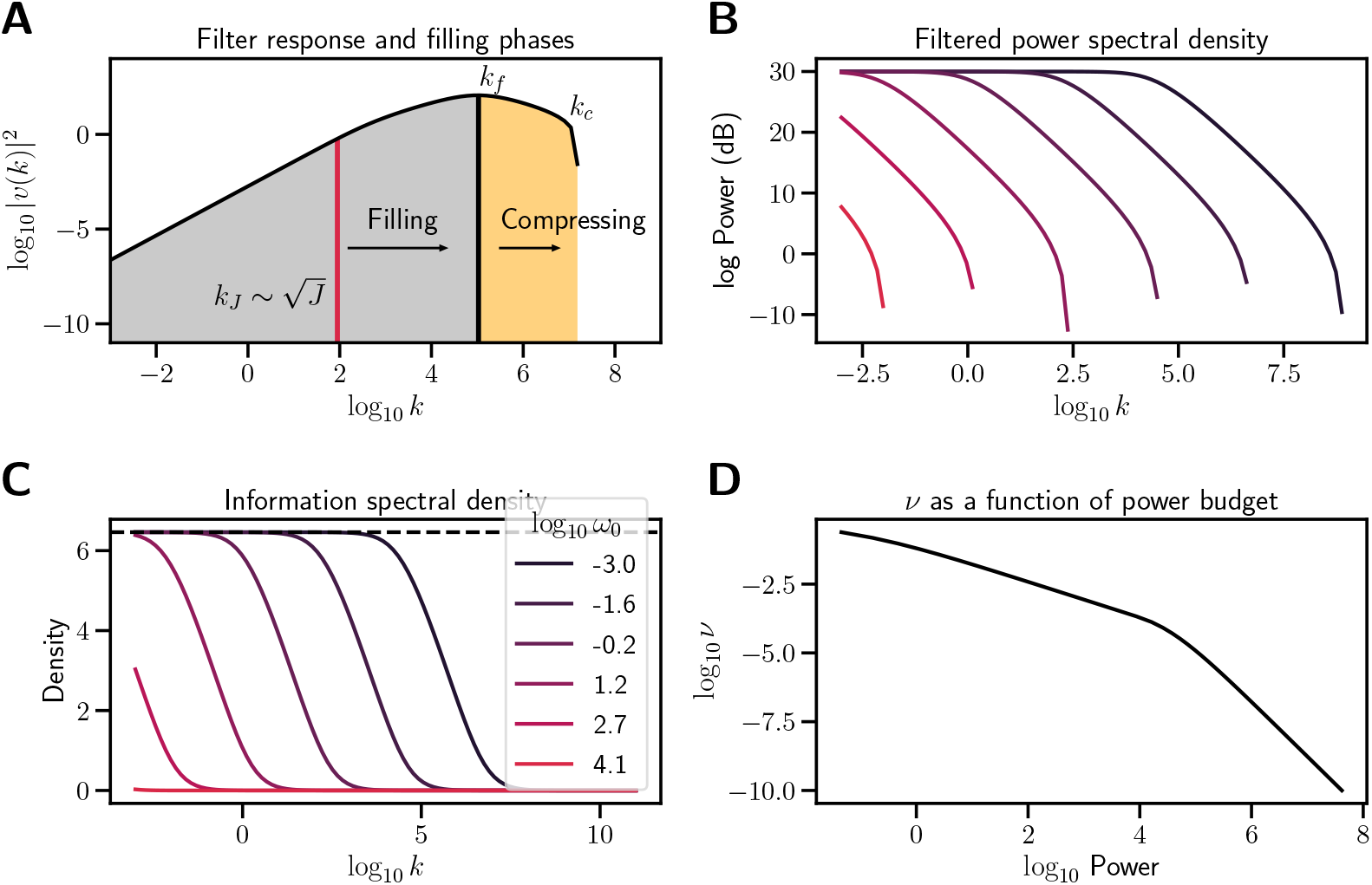
Information as a function of bandwidth. **(A)** Schematic of the optimal filter. The optimal filter response comprises a log-linear region for *k* < *k_f_* and a negative log-linear region for *k_f_* < *k* < *k_c_*. As RGC number *J* ~ *k*^2^ increases through the former, the mosaic fills, pushing the bandwidth limit (red line) upward. When the mosaic is filled, the upper bandwidth limit continues to increase as new RFs pack into the fixed retinal space and compress the mosaic. **(B)** Optimal filter power spectral density for different values of characteristic temporal response frequency *ω*_0_. Brighter colors indicate increasing frequency (see legend in **C**). In each case, the filtered spectrum is flat and proportional to *ν*^-1^ for *k* < *k_f_*. For *k* > *k_f_*, the power is noise-dominated, with a *k*^-α/2^ tail. **(C)** Information as a function of frequency for the same setup as in **B**. Note that information content falls precipitously for *k* > *k_f_* (the y axis is not logged). The dotted line indicates – 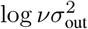. **(D)** As the power budget increases, ν decreases, with ν ~ P^-1^ at low power and *ν* ~ *P*^-2^ as *P* → ∞. For all plots, *A* = 100, σ_in_ = 0.4, σ_out_ = 1.25. In **A–C**, *ν* = 10^-3^. In **A**, log_10_ *ω*_0_ = −2.3; in **D**, log_10_ *ω*_0_ = 1.2.

We can gain additional insight by examining two important regimes in this distribution: If *k* ≳ *k_f_* then

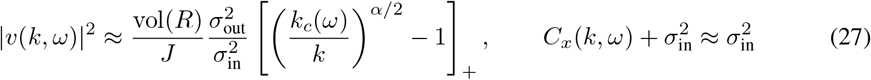

consistent with the definition of *k_c_* as an *ω*-dependent cutoff frequency. Conversely, when *k* ≲ *k_f_* (*ω*),

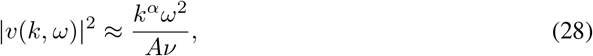

independent of *J*, the filter is separable, and the image spectrum is white [9]:

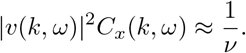

Supplementary Figure 1A shows that indeed, the log-scaled filter magnitude is characterized by a linear regime when *k* ≲ *k_f_*, followed by a 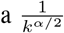 decay when *k* ≳ *k_f_*. Thus, despite the fact that power density is highest in the first regime, the second is light-tailed and stretches exponentially longer, with the result that total power is dominated by the tail (Supplementary Figure 1B). And indeed, numerical integration shows that power initially scales as *ν*^-1^ for large values, followed by *P* ~ *ν*^-1/2^ as *ν* → 0, consistent with this claim (Supplementary Figure 1D). By contrast, as Supplementary Figure 1C shows, information density (log terms in (20)) falls off much more rapidly with frequency, such that it is the *k* ≲ *k_f_* regime that dominates, and information 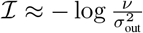.

However, the above analysis is all in the continuous (infinite retina case). For the finite retina case, we note two important changes: First, as the number of RGCs increases, so does the power budgeted to the system. That is, the allowed power is assumed to be *P*(*J*) = *P*_0_*J*. This need not be the case — some other scaling could just as well be chosen — but it is consistent with the idea that metabolic costs are driven by factors specific to each cell. Second, the information in the system is no longer given by the the first term in the integral (20) but by a sum over discrete frequencies 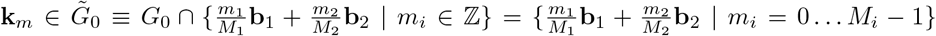. For example, when *J* =1, *M*_1_ = *M*_2_ = 1 and *m*_1_ = *m*_2_ = 0 is the only option — the periodicity of the signals is the periodicity of the lattice — whereas *J* = 2 allows either *M*_1_ =2 or *M*_2_ = 2 but not both, and *J* = 4 has *M*_1_ = *M*_2_ = 2. In general, for large enough *J*, we expect 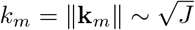. With these conventions, we then have modified expressions for both the information in the system and its power consumption:

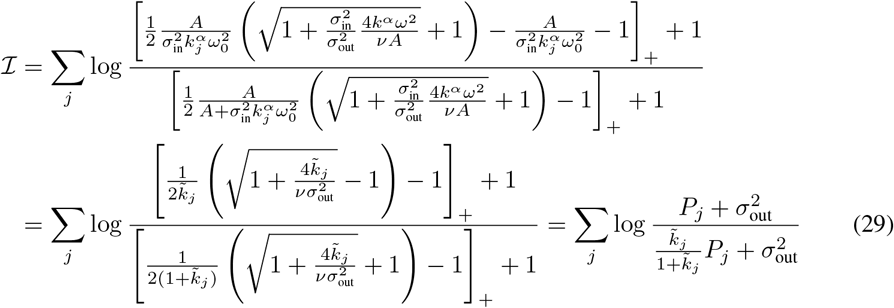

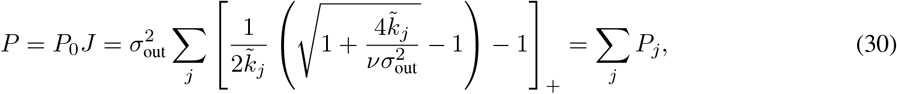

where we have chosen to focus on filters narrowly concentrated around a single frequency *ω*_0_ and again used 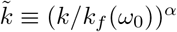.

Several things are important to note about the scaling relationships in (29) and (30). First, as discussed above, for 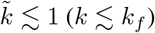,

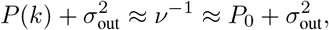

independent of *k*. That is, contributions to (30) are all roughly the same for each frequency. Similarly, the denominator in (29) is 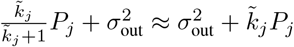 to lowest order in 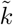, giving

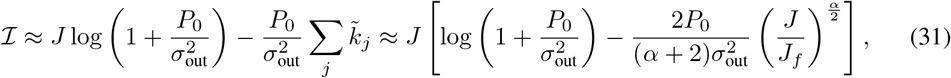

where we have used the volume equivalence 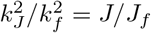 along with the integral approximation

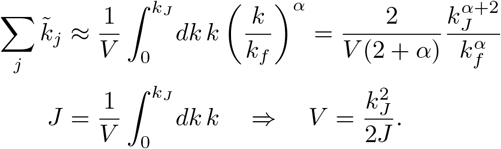

We can understand this *k* ≲ *k_f_* regime as a *mosaic filling phase*: For small energy budgets 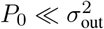, new RFs are added at their preferred size 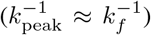 until the spacing between RFs reaches a minimum of roughly 2*π*/*k_f_* (at which point 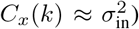), with information increasing almost linearly as the mosaic covers new spatial locations. By contrast, for larger energy budgets 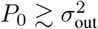, information gain is sublinear in *J* with a correction term ~ *J*^*α*/2^.

By contrast, for *k* ≳ *k_f_*, and the system enters a *mosaic compression phase*. In this regime, noise dominates signal, 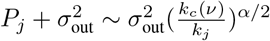, and for the power above *k_f_*, we have

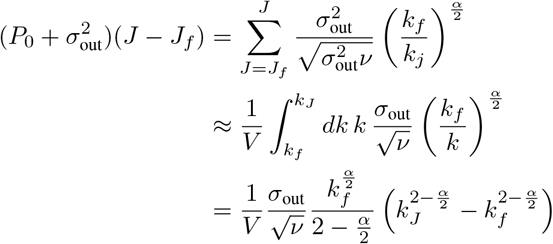

where *J_f_* is the number of RGCs contained in *k* ≲ *k_f_* and *V* is once again a normalizing factor required to convert the sum over discrete frequencies into the integral over *k*:

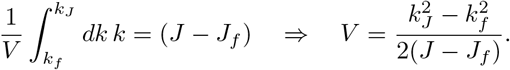

Putting these together and using 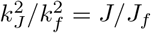 in two dimensions then gives

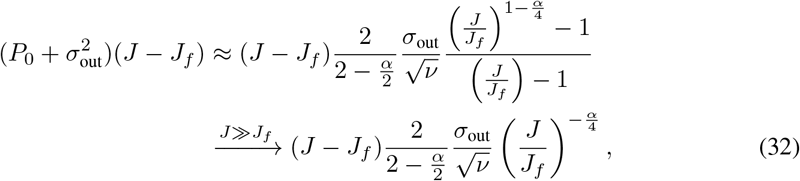

which requires *ν* ~ (*J*/*J_f_*)^*α*/2^ asymptotically for proper power scaling.

From (29), the compression phase adds then adds information

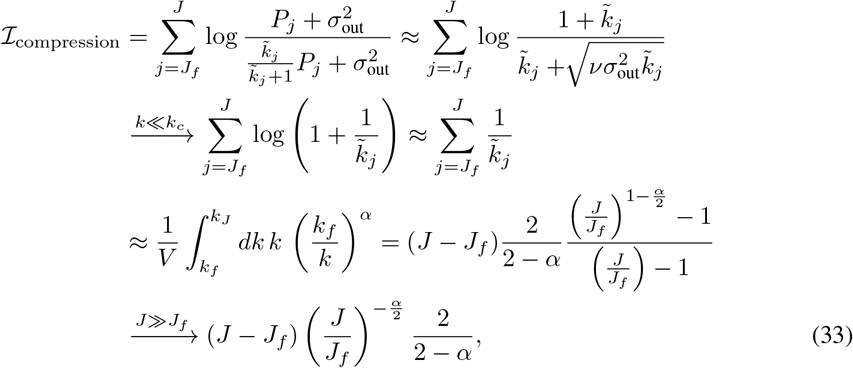

which is again sublinear growth in *J*, but not so strong as the decay in (31) when 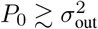.

Finally, we consider the effects of temporal frequency *ω* on the relationships we have described above. In the case we have assumed, that of a filter with a narrow temporal passband centered around *ω*_0_, we note that (29) depends only on 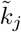, and we have

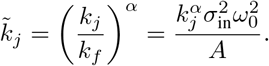

Thus, if we increase *ω*_0_ → *θω*_0_ with *θ* > 1, we have 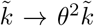 which moves the numerator and denominator in (29) closer to each other and information to 0. The effect of this change in temporal frequency can be seen in Figure 2E. As a result, mosaics with the lowest temporal frequency are filled first, with new mosaics only beginning to fill once the marginal information gain for adding the first RF to a new mosaic exceeds that of adding an additional RF to an existing mosaic. In this way, the rate of each mosaic’s filling decreases as new mosaics are added Figure 2F.

**Supplementary Figure 2:**
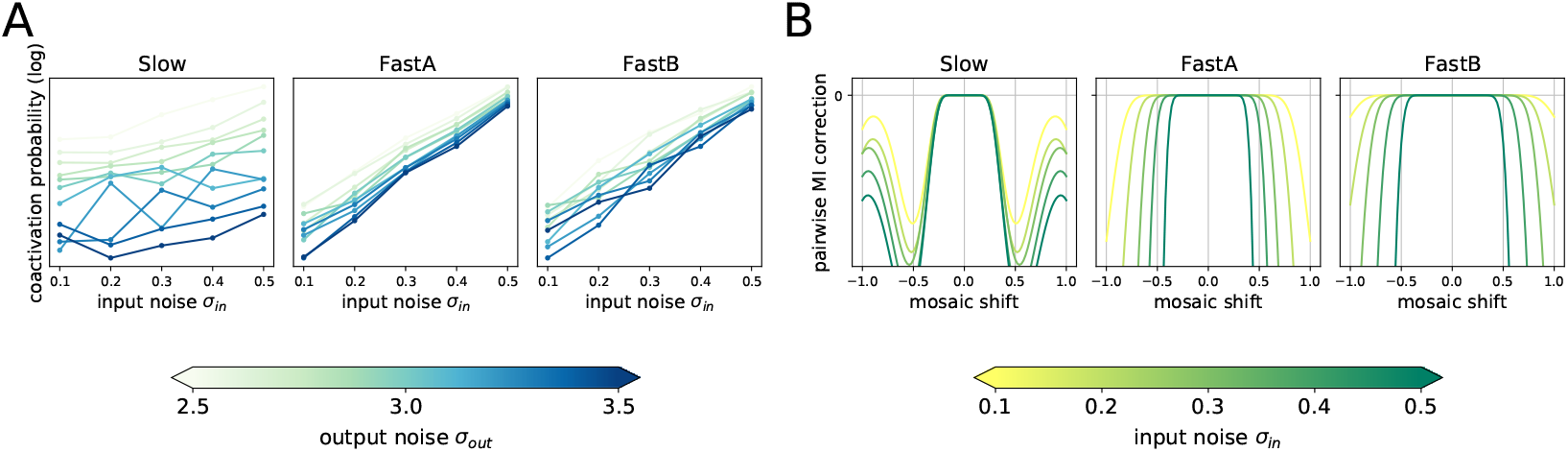
**(A)** Across all output noise levels, the coactivation probability of an ON RGC at the center and an OFF RGC at the edge increases as the input noise σ_in_ increases. **(B)** The pairwise mutual information correction term, which captures the reduction in mutual information due to pairwise coactivations of RFs (plotted for σ_out_ = 3.0) stays near zero for a shorter range of mosaic shifts at higher input noise levels, indicating that higher input noise favors aligned mosaics.

## B Analysis of mosaic phase transitions as a function of input noise

Figure 5 in the main text shows that, for each filter type, increasing input noise levels encourage alignment. That is, the transition happens at a higher output noise level. Here, we explain this seemingly paradoxical effect using the mathematical analysis presented in [14], whose notation we follow. In that work, it was argued that the transition from aligned to anti-aligned mosaics could be understood as a process of increasing output noise leading to higher response thresholds, which in turn decorrelated nearby ON and OFF cells [12, 14]. As these pairs become more independent (as measured by *p*_2_, their pairwise coactivation probability), they remain so even for small relative shifts in position, implying that anti-alignment does not increase coding redundancy. Add to this the fact that anti-aligned mosaics sample more unique spatial positions, and the result is that high output noise levels lead to anti-alignment under efficient coding.

Conversely, though not analyzed in [14], the observation that increasing input noise levels favor alignment can be explained through the same formalism. Whereas [14] found that, in the independent neuron case, increasing output noise drove increases in response threshold (to better encode stimuli above the noise floor), the same analysis shows that increases in input noise *decrease* response thresholds (Supplementary Figure 3). Moreover, [14] showed that the coactivation probability between an ON-OFF RF pair at a distance of *j* lattice positions away in *d* = 1 with Gaussian inputs is given by

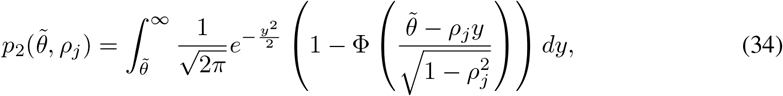

where, for 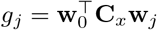 and 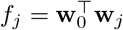,

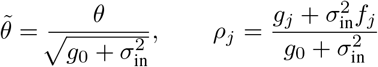

are the signal-to-noise-normalized threshold and unthresholded ON-OFF correlation, respectively. From this, it is clear (and Supplementary Figure 2A shows) that increasing *σ*_in_ reduces 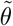 not only through reductions in the optimal *θ*, but through increases in the normalization factor. This, in turn, reduces *p*_2_ through an increase in the integration limits in (34). Thus, increasing *σ*_in_ results in a *narrower* “zone of independence” (Supplementary Figure 2B) around each RF, with the result that mosaics cannot anti-align without losing information.

## C Sinusoidal temporal kernels in the absence of parameterization

Supplementary Figure 5 shows the distribution of temporal filters learned without using the parameterization in Equation 6, which have symmetric, sinusoidal shapes resembling Fourier bases. They also exhibit distinct clusters of temporal kernel shapes with varying spectral characteristics, and each group corresponds to a spatial RF mosaic that tiles the space (Supplementary Figure 5E). Supplementary Figure 5A-D shows that the temporal filters are almost perfectly orthogonal across the groups, which allows independent information to be conveyed in each group. The shapes of the temporal filters obtained in the full model (Supplementary Figure 3 and 4) can be interpreted as being as independent as possible while retaining biologically plausible filter shapes. In the case of auditory filters, it has been shown that efficient coding results in Fourier-like filters [27].

**Supplementary Figure 3:**
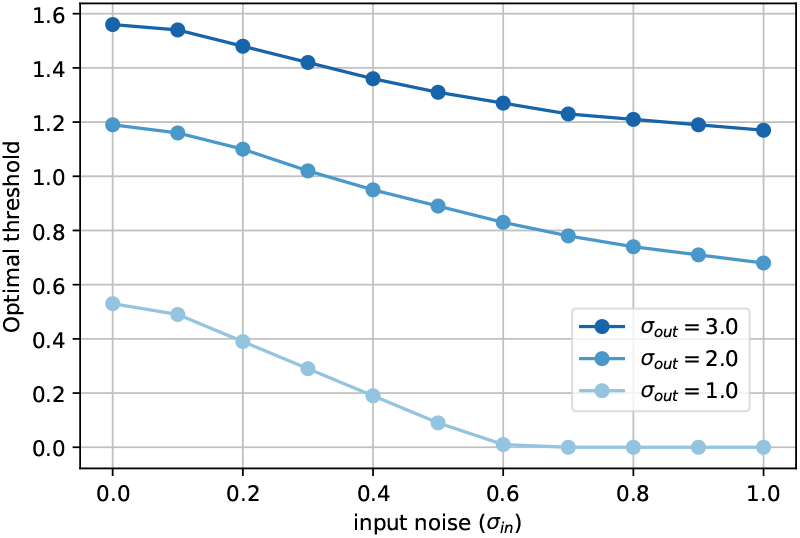
Optimal thresholds decrease with input noise as they increase with output noise. Calculations based on the independent neuron model of [14].

## D Information gain upon adding individual RF mosaics

We examine how each of the RF mosaics contribute to the overall mutual information, by manually constructing four pairs of mosaics of spatial kernels placed in hexagonal grids, corresponding to four types of temporal filters: *Slow, FastA, FastB*, and *FastC* (6A). Given the 8 groups of spatiotemporal kernels, Figure 6B shows the order of adding mosaics such that each step brings the largest increase in the estimated mutual information. The optimal ordering indicates that it is always beneficial to add the slow mosaics first, in either order between ON and OFF (steps 1 and 2). Afterwards, the highest increase in MI is when adding one mosaic, either ON or OFF, from each shape in the order of increasing speed (steps 3-5), and then filling in the remaining mosaics again in the order of increasing speed (steps 6-8).

## E Additional Technical Details

### Determining the polarity of spatiotemporal kernels

We used the parameterization of temporal kernels in Equation 6, and we additionally experimented with temporal kernels that can learn any unit-norm function in Supplementary Figure 5. Unlike that of spatial kernels in Equation 8 which has a positive peak intensity at the center, both parameterizations for temporal kernels are symmetric as to the positive and negative values the kernels can take, and there is no built-in indicator to distinguish ON and OFF kernels. Since ON and OFF RFs are the ones responding, respectively, to light increment and decrement [28], we determine the polarity of given temporal kernel *w*[*t*] based on the sign of its cumulative response to the unit step function *u*[*t*] up to the half duration of the kernel:

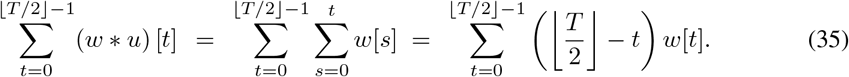

In the visualizations of the spatial and temporal kernels, we flip the signs of the OFF temporal kernels as well as the corresponding spatial kernels, which keeps the separable spatiotemporal kernels unchanged (Equation 5). This makes the temporal kernels always visualized as ON filters, and the spatial kernels are displayed with a bright center for ON RFs and a dark center for OFF RFs. We also note that we plot the temporal kernels so that *t* = 0 is at the right and *T* = *t* - 1 is at the left of the graphs, making it comparable to the usual direction of visualizing temporal RFs, e.g. in spike-triggered averages.

**Supplementary Figure 4:**
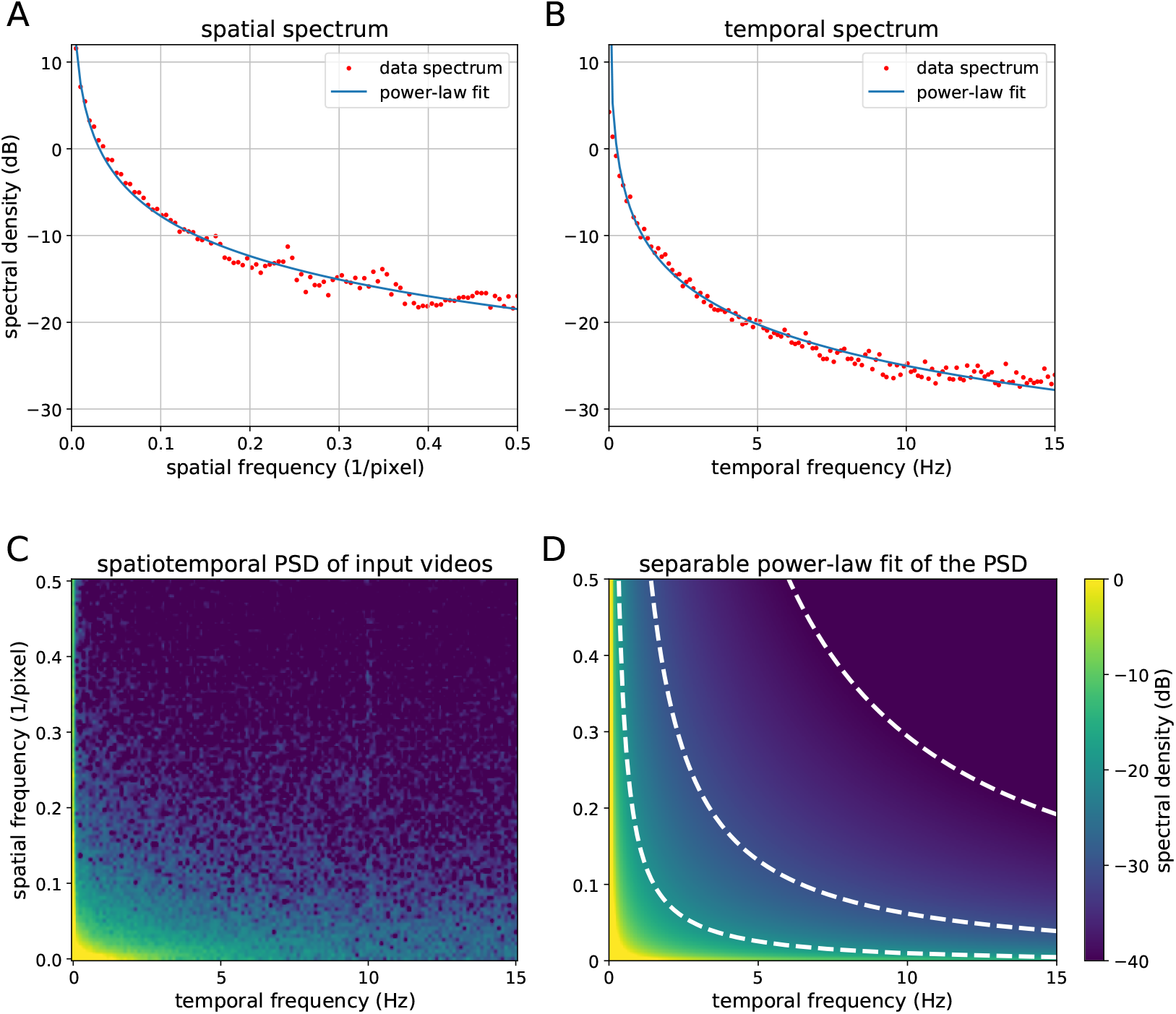
The spatiotemporal spectrum of the input videos from the Chicago Motion Database is well-approximated by separable power-law fits of the spatial and temporal spectra. **(A)** The spatial spectrum of the data and its power-law fit. **(B)** The temporal spectrum of the data and its power-law fit. **(C)** The spatiotemporal power-spectral density of the data. **(D)** Reconstructed power-spectral density from the power-law fits. Contour lines indicate −40 dB, −30 dB, and −20 dB.

### Data preprocessing and training

The Chicago Motion Database [22] contains 257 natural video clips ranging from bees flying around the hive to waves approaching the shore. All video clips in the dataset are square-shaped and have varying sample rates. We resize and resample the video clips to 512×512 pixels and 30 fps using ffmpeg, in addition to converting it to grayscale by taking the luminosity channel of each frame using the Python Imaging Library. Following [10] and [14], we normalize each video to have zero mean and unit variance over all video dimensions, while the sliced video patches may have different means and variances. We used square-shaped input patches **x**(*t*) of size *D* = 18^2^ = 324 pixels unless otherwise specified, and we applied a circular mask on the input patches following [14], which effectively constrained the spatial kernels within a circle bounded by the input square. Input patches consisted of either *T* =10 or 20 frames sliced from 30-fps videos, and for computational efficiency, we used “valid”-mode convolutions with temporal kernels of the same size T, so each RGC yielded a single firing rate value *r_j_* for each input. Models using larger convolutions, capturing information across both RGCs and time, produced similar results. For computational efficiency in large-scale experiments in Sections 4.1 and 4.2, we approximated the input data distribution as a multivariate Gaussian matched to the statistics of the natural image dataset estimated using one million samples of video patches. A batch of 128 such video patches was used for each update, and we used Adam optimizer [23] with a constant learning rate of 0.001. In Section 4.1, a model optimizing J RGCs is trained for 5000J iterations, and each model in Section 4.2 is trained for 200,000 iterations. The experiments in 4.2 often result in either of aligned and anti-aligned configurations depending on the initialization of the model, so we repeat each configuration 10 times and select the result that obtained the highest mutual information objective (Equation 2). Each experiment in Sections 4.1, and 4.2 were run on a NVIDIA 1080 Ti GPU for about 50 hours, 20 hours, and 5 hours, respectively.

**Supplementary Figure 5:**
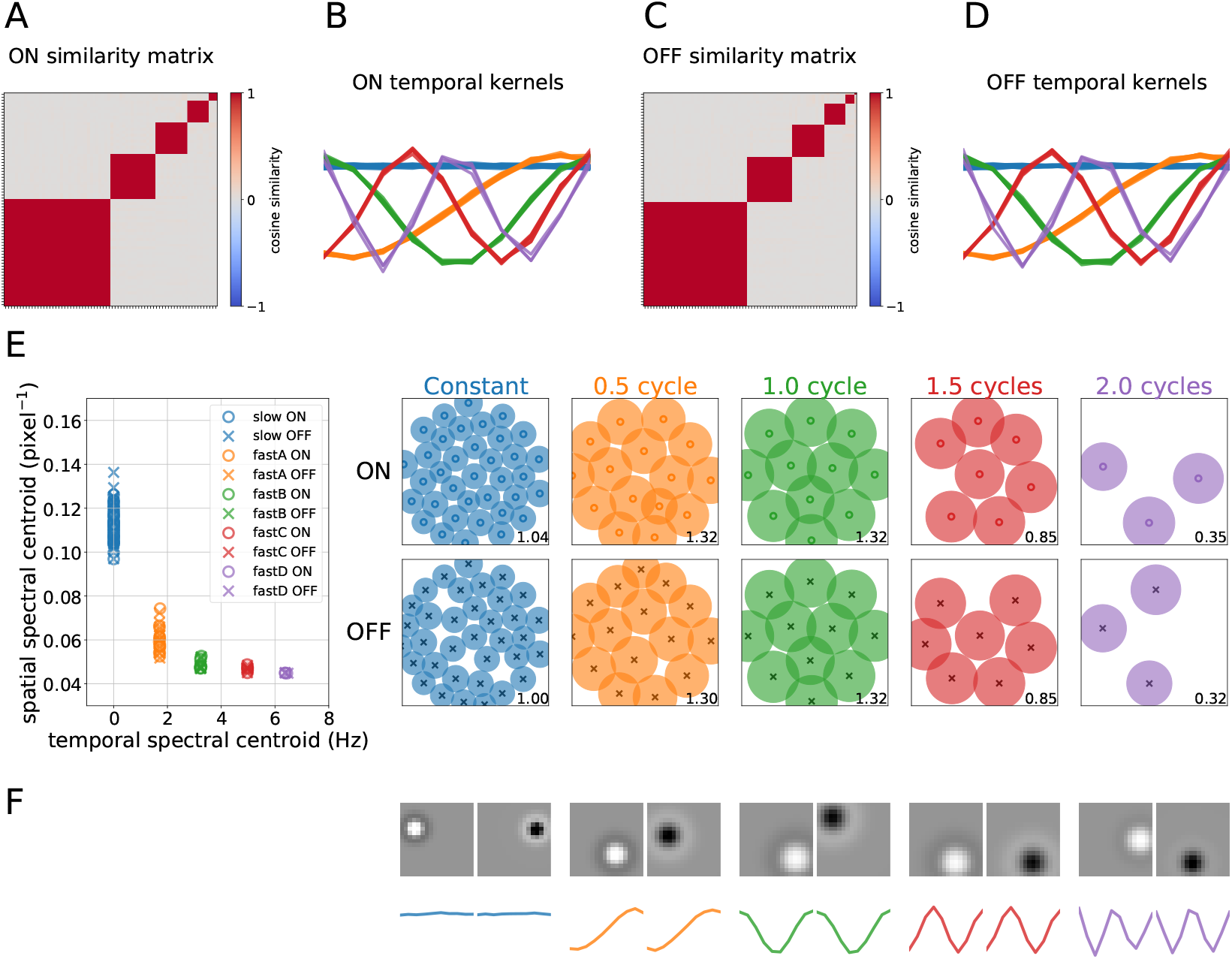
Sinusoidal bases are learned when any unit-norm temporal kernels of a given length are allowed instead of using the parameterization in Equation 6. A model with *J* = 144 is used. **(A, C)** Self-similarity matrices of ON (A) and OFF (C) temporal filters show five distinct clutsers in each, with almost perfect orthogonality across clusters. The filters are sorted according to their spectral centroids. **(B, D)** The shapes of learned temporal ON (B) and OFF (D) filters, color-coded according to the clusters. Each cluster exhibits a sinusoidal shape with a distinct frequency, ranging from the constant to having roughly 0.5, 1.0, 1.5, and 2.0 cycles in the given kernel size. **(E)** (left) The distribution of the temporal and spatial spectral centroids. (right) ON and OFF mosaics corresponding to each RF type. The number in the corner of each plot denotes the coverage factor of the mosaic. **(F)** Learned shapes of a spatial ON and OFF filters (top) and the corresponding temporal filters (bottom) from each RF type.

**Supplementary Figure 6:**
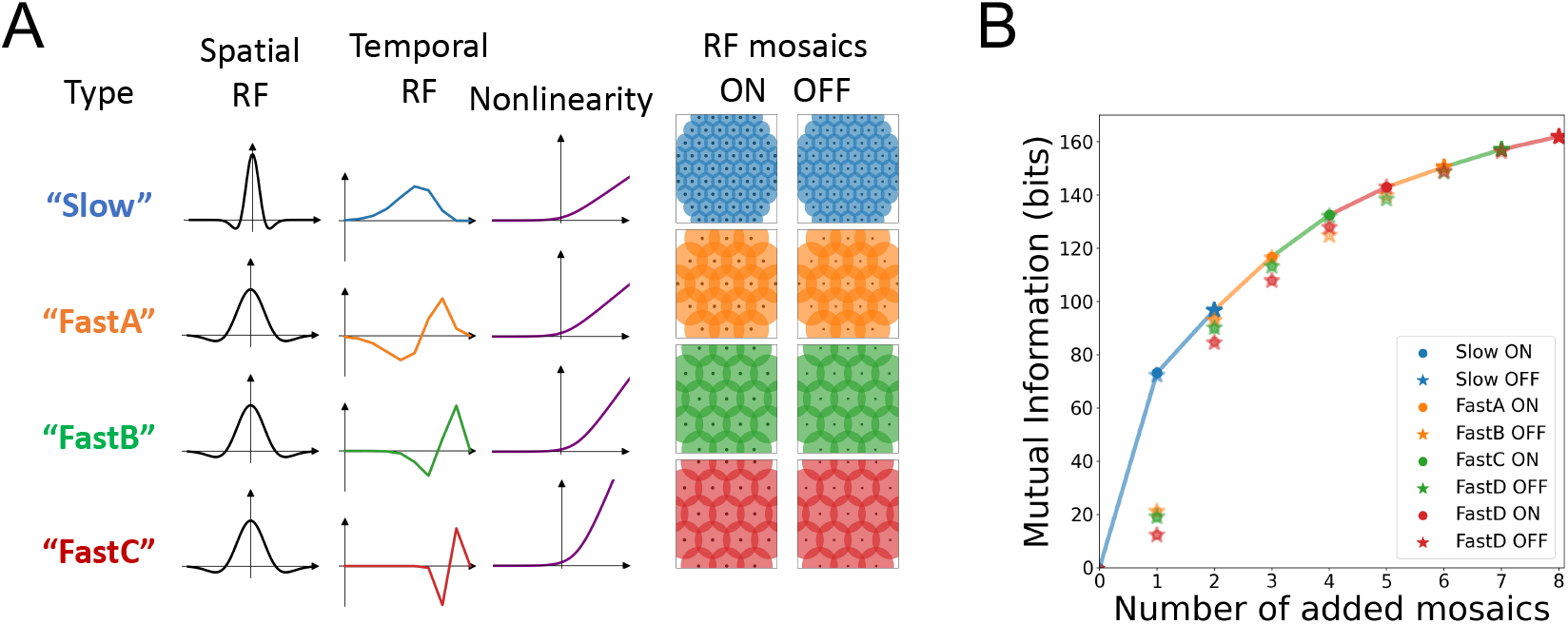
The order of mosaic emergence is explained by the information gain upon adding individual mosaics. **(A)** Four sets of parameters are manually selected to construct spatial and temporal RFs, nonlinearities, and RF mosaics that resemble the *Slow, FastA, FastB*, and *FastC* RGCs. The mosaics are constructed to pack the space using a hexagonal pattern, resulting in *J* = 234 RGCs across the 8 mosaics. **(B)** Starting from an empty set, we repeat finding and adding the mosaic that maximally increases the mutual information, until all 8 mosaics are included. At each step, a scatter plot of the mutual information for each choice is plotted, and the solid line indicates the increase in mutual information by adding the best mosaic at each step.

**Supplementary Figure 7:**
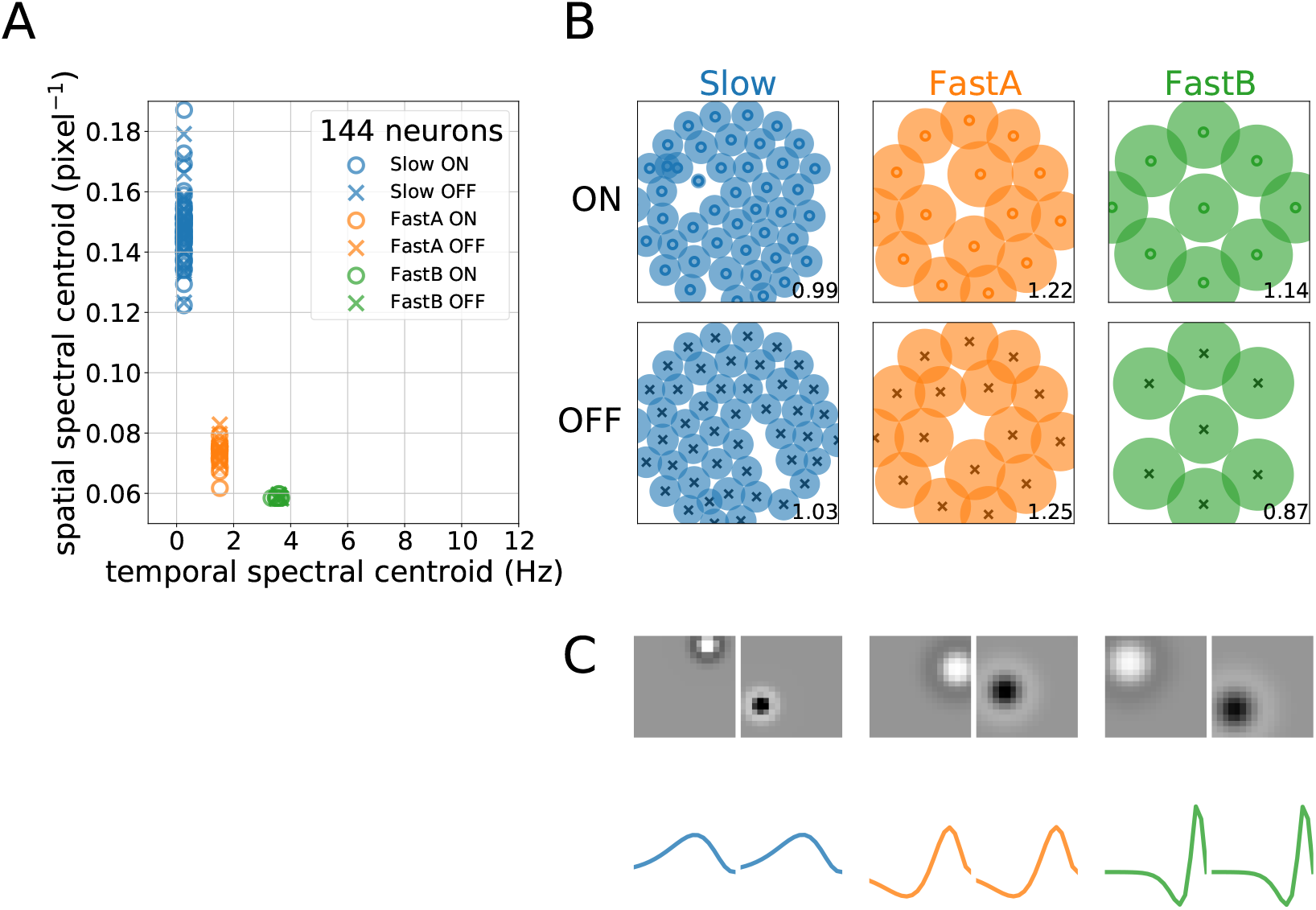
Emergence of RF mosaics is not contingent on the choice of the hyperparameter n in Equation 6. The above is the result of optimizing *J* = 144 RGCs using *n* = 3 instead of the default n = 6, showing the resulting distribution of spatiotemporal spectral centroids (A) and RF mosaics (B), as well as their spatial and temporal shapes (C). While the exact trends are different from the case of *n* = 6, this result indicates that the gradual emergence of distinct RF mosaics with increasing J would happen with different values of *n*.

**Supplementary Figure 8:**
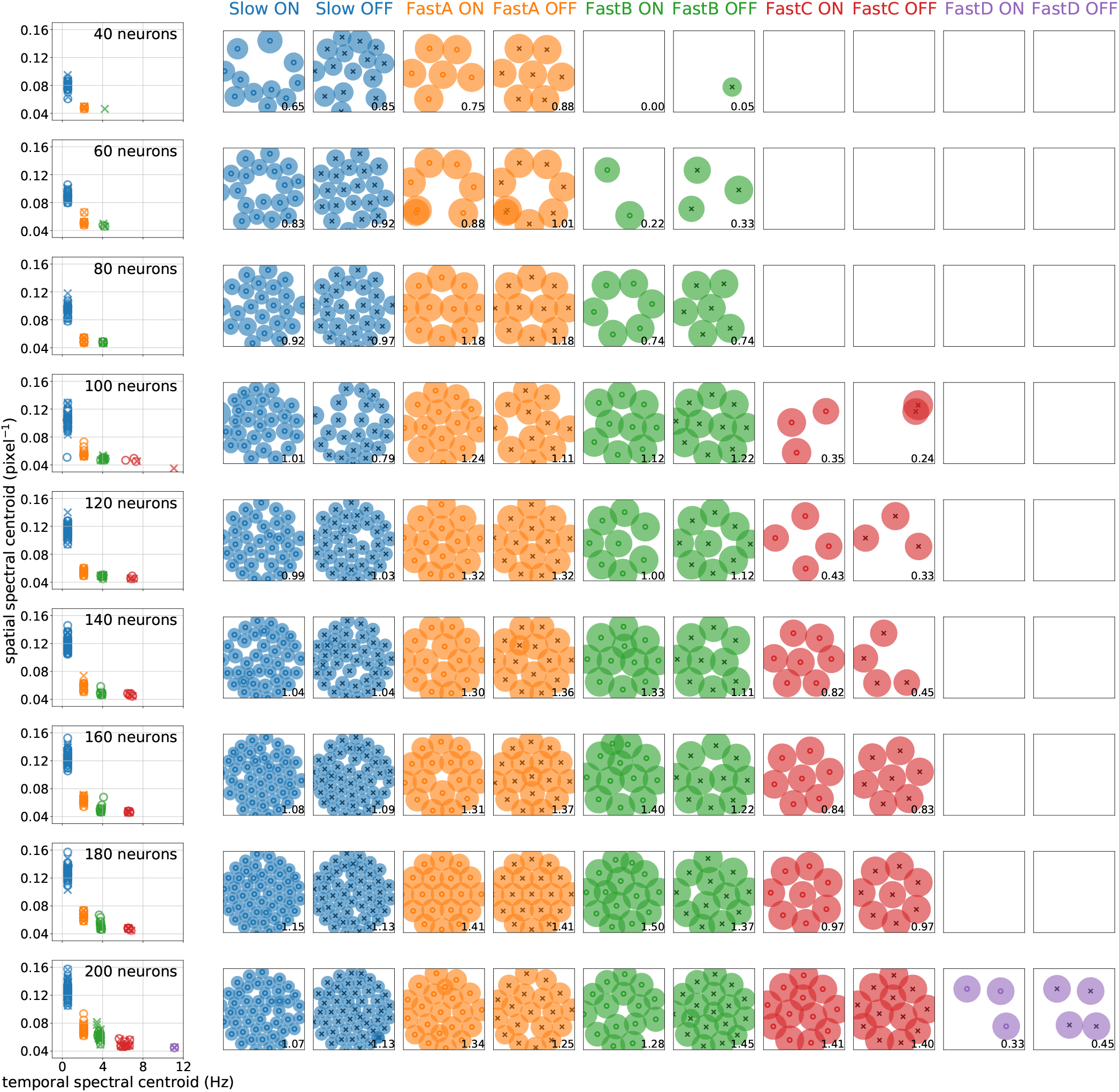
The distribution of the spatial and temporal centroids of kernels (left), the RF mosaics (middle), and the temporal kernel shapes (right), as the number of RGCs under optimization increases from 40 to 200, by an increment of 20. As the number of RGCs increases, faster mosaics emerge to cover the higher-frequency region of the spatiotemporal spectrum, while the individual spatial RFs become more precise, using more number of RGCs to tile the same visual space.

**Supplementary Figure 9:**
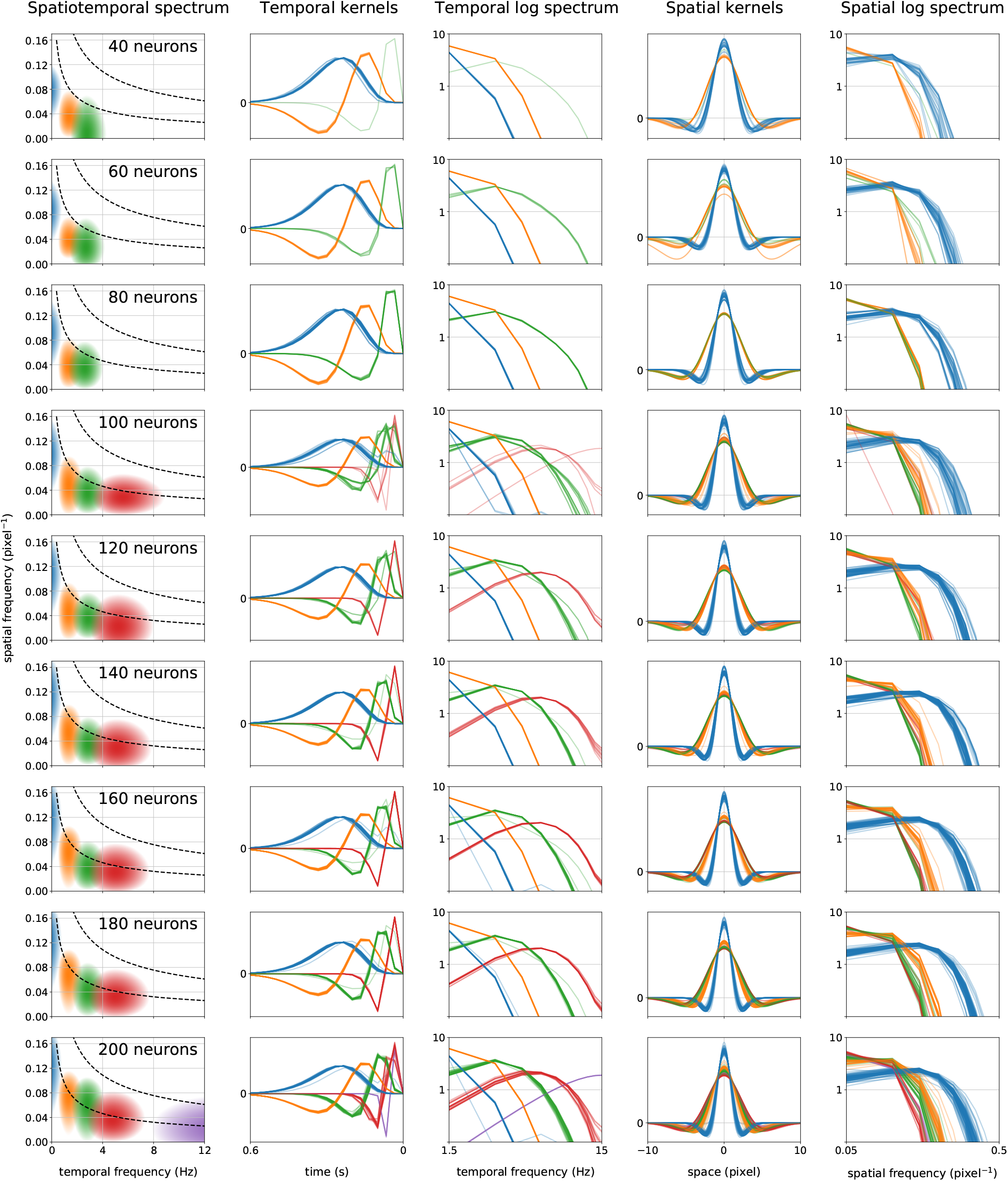
More visualizations of the kernels in Supplementary Figure 8. Column 1 plots the region where the average spatiotemporal kernel of each type has high frequency response, superimposed with contour lines of the dataset’s power spectral density. Each kernel type covers a different region of spatiotemporal frequencies, together packing increasingly larger areas where the spectral power of the dataset is concentrated. Individual temporal kernels are plotted in the time and frequency domains (columns 2-3), and spatial kernels in the space and frequency domains (columns 4-5).

**Supplementary Figure 10:**
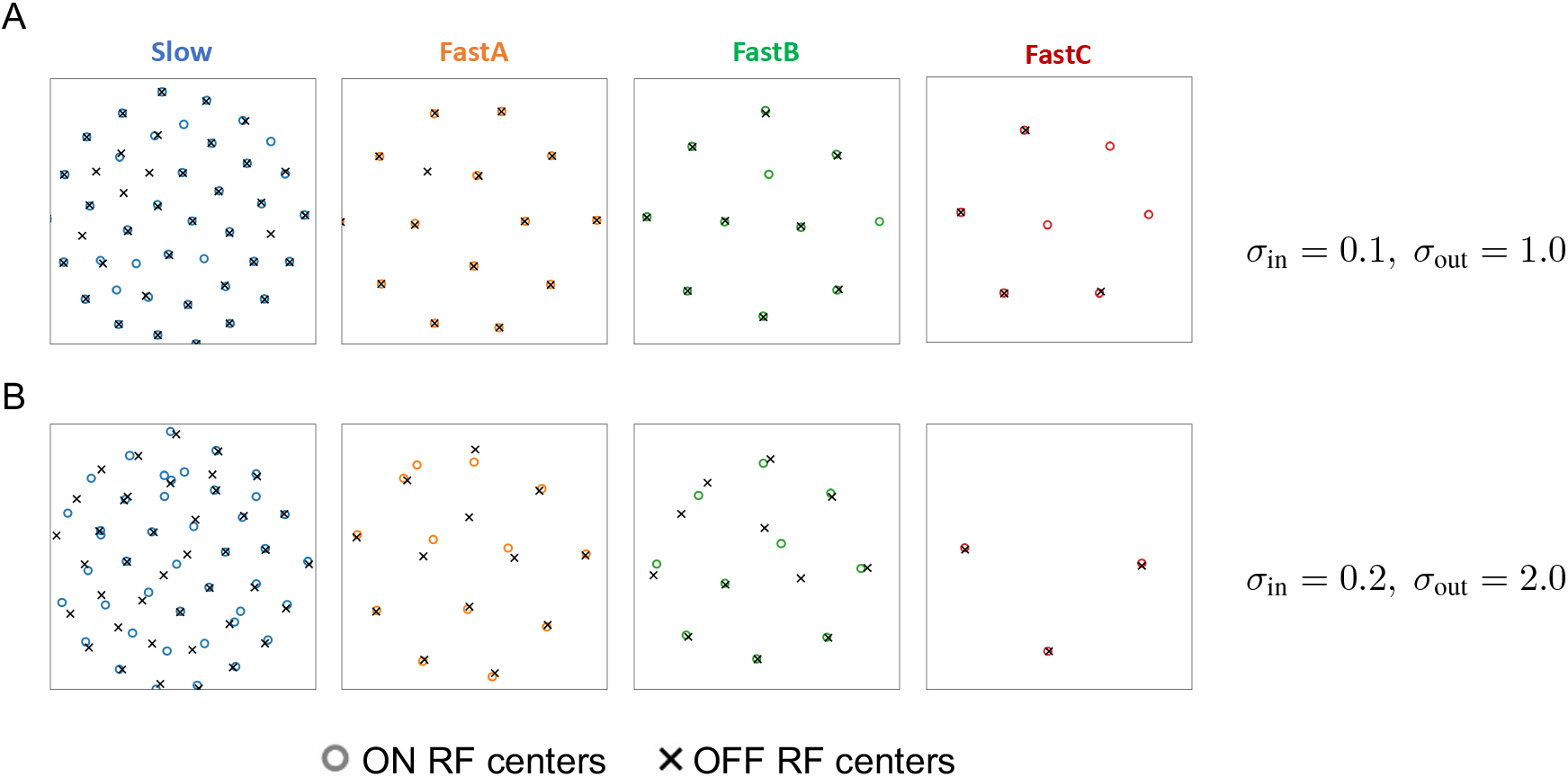
RF centers per mosaic type under different input and output noise levels, using *J* = 140 RGCs. ON and OFF RF centers are more aligned under a lower noise level than under a higher noise level.

## Notes

https://github.com/pearsonlab/efficientcoding

### Competing Interest Statement

The authors have declared no competing interest.

### Summary of Updates

removed a typo in the abstract

## References

[1] Mrinalini Hoon, Haruhisa Okawa, Luca Della Santina, and Rachel OL Wong. Functional architecture of the retina: development and disease. Progress in retinal and eye research, 42:44–84, 2014.

[2] H Wässle, L Peichl, and Brian Blundell Boycott. Morphology and topography of on-and off-alpha cells in the cat retina. Proceedings of the Royal Society of London. Series B. Biological Sciences, 212(1187):157–175, 1981.

[3] Steven H Devries and Denis A Baylor. Mosaic arrangement of ganglion cell receptive fields in rabbit retina. Journal of neurophysiology, 78(4):2048–2060, 1997.

[4] Greg D Field, EJ Chichilnisky, et al. Information processing in the primate retina: circuitry and coding. Annual review of neuroscience, 30(1):1–30, 2007.

[5] Joshua R Sanes and Richard H Masland. The types of retinal ganglion cells: current status and implications for neuronal classification. Annual review of neuroscience, 38:221–246, 2015.

[6] Fred Attneave. Some informational aspects of visual perception. Psychological review, 61(3):183, 1954.

[7] Horace B Barlow. Possible principles underlying the transformation of sensory messages. Sensory communication, 1:217–234, 1961.

[8] Joseph J Atick and A Norman Redlich. Towards a theory of early visual processing. Neural computation, 2(3):308–320, 1990.

[9] Joseph J Atick and A Norman Redlich. What does the retina know about natural scenes? Neural computation, 4(2):196–210, 1992.

[10] Yan Karklin and Eero P Simoncelli. Efficient coding of natural images with a population of noisy linear-nonlinear neurons. In Advances in neural information processing systems, pages 999–1007, 2011.

[11] Eizaburo Doi, Jeffrey L Gauthier, Greg D Field, Jonathon Shlens, Alexander Sher, Martin Greschner, Timothy A Machado, Lauren H Jepson, Keith Mathieson, Deborah E Gunning, et al. Efficient coding of spatial information in the primate retina. Journal of Neuroscience, 32(46):16256–16264, 2012.

[12] Xaq Pitkow and Markus Meister. Decorrelation and efficient coding by retinal ganglion cells. Nature neuroscience, 15(4):628–635, 2012.

[13] Samuel Ocko, Jack Lindsey, Surya Ganguli, and Stephane Deny. The emergence of multiple retinal cell types through efficient coding of natural movies. In Advances in Neural Information Processing Systems, pages 9389–9400, 2018.

[14] Na Young Jun, Greg D Field, and John Pearson. Scene statistics and noise determine the relative arrangement of receptive field mosaics. Proceedings of the National Academy of Sciences, 118(39), 2021.

[15] Suva Roy, Na Young Jun, Emily L Davis, John Pearson, and Greg D Field. Inter-mosaic coordination of retinal receptive fields. Nature, 592(7854):409–413, 2021.

[16] Audrey J Sederberg, Jason N MacLean, and Stephanie E Palmer. Learning to make external sensory stimulus predictions using internal correlations in populations of neurons. Proceedings of the National Academy of Sciences, 115(5):1105–1110, 2018.

[17] EJ Chichilnisky and Rachel S Kalmar. Functional asymmetries in on and off ganglion cells of primate retina. Journal of Neuroscience, 22(7):2737–2747, 2002.

[18] Tom Baden, Philipp Berens, Katrin Franke, Miroslav Román Rosón, Matthias Bethge, and Thomas Euler. The functional diversity of retinal ganglion cells in the mouse. Nature, 529(7586):345–350, 2016.

[19] Sneha Ravi, Daniel Ahn, Martin Greschner, EJ Chichilnisky, and Greg D Field. Pathway-specific asymmetries between on and off visual signals. Journal of Neuroscience, 38(45):9728–9740, 2018.

[20] W. Bialek and W.G. Owen. Temporal filtering in retinal bipolar cells. elements of an optimal computation? Biophysical Journal, 58(5):1227–1233, 1990.

[21] Dawei W Dong and Joseph J Atick. Statistics of natural time-varying images. Network: Computation in Neural Systems, 6(3):345, 1995.

[22] Jared M. Salisbury and Stephanie E. Palmer. Optimal prediction in the retina and natural motion statistics. Journal of Statistical Physics, 162(5):1309–1323, January 2016.

[23] Diederik P Kingma and Jimmy Ba. Adam: A method for stochastic optimization. arXiv preprint arXiv:1412.6980, 2014.

[24] Jorge Nocedal and Stephen Wright. Numerical optimization. Springer Science, 35(67-68):7, 1999.

[25] Stephanie E Palmer, Olivier Marre, Michael J Berry, and William Bialek. Predictive information in a sensory population. Proceedings of the National Academy of Sciences, 112(22):6908–6913, 2015.

[26] Alan V Oppenheim, Ronald W Schafer, Mark A Yoder, and Wayne T Padgett. Discrete-time signal processing. Pearson, Upper Saddle River, NJ, 3 edition, August 2009.

[27] Michael S Lewicki. Efficient coding of natural sounds. Nature neuroscience, 5(4):356–363, 2002.

[28] Stephen W Kuffler. Discharge patterns and functional organization of mammalian retina. Journal of neurophysiology, 16(1):37–68, 1953.

